# Transferability of cathodal tDCS effects from the primary motor to the dorsolateral prefrontal cortex: a multimodal TMS-EEG study

**DOI:** 10.1101/2022.06.13.495748

**Authors:** Mohsen Mosayebi-Samani, Desmond Agboada, Tuomas P. Mutanen, Jens Haueisen, Min-Fang Kuo, Michael A. Nitsche

## Abstract

Neurophysiological effects of transcranial direct current stimulation (tDCS) have been extensively studied over the primary motor cortex (M1). Much less is however known about its effects over non-motor areas, such as the prefrontal cortex (PFC), which is the neuronal foundation for many high-level cognitive functions and involved in neuropsychiatric disorders. In this study, we, therefore, explored the transferability of cathodal tDCS effects over M1 to the PFC. Eighteen healthy human participants (11 males and 8 females) were involved in eight randomized sessions, in which four cathodal tDCS dosages, low, medium, and high, as well as sham stimulation, were applied over the M1 and PFC. After-effects of tDCS were evaluated via transcranial magnetic stimulation (TMS)-electroencephalography (EEG), and TMS-elicited motor evoked potentials (MEP), for the outcome parameters TMS-evoked potentials (TEP), TMS-evoked oscillations, and MEP amplitude alterations. TEPs were studied both at the regional and global scalp levels. The results indicate a regional dosage-dependent nonlinear neurophysiological effect of M1 tDCS, which is not one-to-one transferable to PFC tDCS. Low and high dosages of M1 tDCS reduced early positive TEP peaks (P30, P60), and MEP amplitudes, while an enhancement was observed for medium dosage M1 tDCS (P30 and MEP amplitudes). In contrast, prefrontal low, medium and high dosage tDCS uniformly reduced the early positive TEP peak amplitudes. Furthermore, for both cortical areas, regional tDCS-induced modulatory effects were not observed for late TEP peaks, nor TMS-evoked oscillations. However, at the global scalp level, widespread effects of tDCS were observed for both, TMS-evoked potentials and oscillations. This study provides the first direct physiological comparison of tDCS effects applied over different brain areas and therefore delivers crucial information for future tDCS applications.

**SIGNIFICANCE STATEMENT:** Modulatory effects of tDCS over the M1 were largely taken as a template so far for the use of this intervention over non-motor regions. However, the neurophysiological effects of tDCS over non-motor regions, such as the prefrontal cortex (PFC), have been much less explored. In the current study, we, using concurrent transcranial magnetic stimulation- electroencephalography, systematically explored the transferability of cathodal tDCS effects on cortical excitability from M1 to the PFC. The results indicate a dosage-dependent nonlinear neurophysiological effect of motor cortex tDCS, which is not one-to-one transferable to prefrontal tDCS. This study provides the first direct physiological comparison of tDCS effects applied over different brain areas, which will further consolidate the rationale for the extension of tDCS applications at both, basic and clinical levels.

## 1. Introduction

Application of weak direct current via electrodes placed over the scalp (transcranial direct current stimulation, tDCS) has been shown to alter cortical excitability over the primary motor cortex (M1) during, but also after the end of the intervention, inducing plasticity-like after-effects. The direction, magnitude, and duration of respective effects depend on stimulation parameters such as polarity and intensity/duration. Here, anodal tDCS, which refers to surface inward current over the target area, within certain dosage limits, typically results in enhancement of motor cortical excitability, whereas cathodal tDCS, which refers to outward current over the target area, reduces it (Nitsche & Paulus, 2000, 2001; Nitsche et al. 2003b). This thus opens a window to shed light on brain functions underlying cognitive functions (Yavari et al., 2018) or alter symptoms of neurological and psychiatric disorders accompanied by pathological alterations of cortical excitability (Lefaucheur et al., 2017).

The neurophysiological effects of tDCS over the M1 were largely taken as a template so far for the use of this intervention over non-motor regions, whereas previous findings show only gradual comparability between the M1 and other cortical areas. Over the sensorimotor cortex, anodal tDCS increased the amplitude of somatosensory potentials, whereas cathodal tDCS had no effects in one study (Matsunaga et al., 2004), while another study showed excitability-diminishing effects of only cathodal tDCS (Dieckhöfer et al., 2006). Over the visual cortex, anodal tDCS enhanced, and cathodal tDCS reduced visual evoked potential amplitudes, however, the duration of the effects was relevantly shorter as compared to M1 stimulation with otherwise identical protocols (Antal et al., 2004). Taking anatomical, as well as receptor and neurotransmitter distribution differences of distinct cortical areas into account, these gradual differences of effects are plausible and require direct physiological tests of tDCS over respective target areas (Yavari et al., 2018).

This is of critical importance, as in addition to the tDCS applications in basic research and clinical settings of the motor domain, its effects have been also extensively explored for the treatment of neuropsychiatric diseases (Kuo et al., 2017), and exploration of physiological mechanisms underlying cognitive functions, with the dorsolateral prefrontal cortex (DLPFC) as target region (Nitsche et al., 2012; Dedoncker et al., 2016; Karuza et al., 2016). However, the neurophysiological effects of tDCS over this area have been much less explored. Application of concurrent transcranial magnetic stimulation (TMS) and electroencephalography (EEG), opened up a window to address the effects of tDCS on cortical excitability of different brain regions, as indexed by TMS-evoked potentials (TEPs) recorded from scalp EEG electrodes (Tremblay et al., 2019).

Few studies have employed TMS-EEG so far to evaluate tDCS effects. ***Over the M1***, anodal enhancement and cathodal inhibition of early TEP amplitudes have been observed for tDCS with 1 mA for 13 min (Pellicciari et al., 2013), and 2 mA for 10 min (Ahn and Fröhlich, 2021). ***For the prefrontal cortex (PFC)***, anodal tDCS with 1 mA for 20 min, applied with bipolar and high-definition tDCS (HD-tDCS), induced likewise an increase of local early TEP peaks, and a decrease of TMS-evoked beta and gamma oscillatory power over posterior EEG channels (Hill et al., 2017, 2019). In another study, with a newly developed electrode configuration, tDCS with 1.5 mA for 14 min, targeting the left DLPFC with the anode, and the right DLPFC with the cathode, however, showed a reduction of late TEPs (about 120 ms after the TMS pulse) over only the parietal cortex, accompanied by a reduction of TMS-evoked power of theta and gamma oscillations at the global scalp level, whereas tDCS with opposite electrode positions had no effects (Gordon et al., 2018b). Thus, most TMS-EEG studies so far indicated qualitatively comparable results of tDCS over the M1 and PFC, however, the number of studies is limited, and results are partially heterogeneous. A systematic comparative investigation of the neurophysiological effects of tDCS over these different brain regions is therefore required.

In a foregoing study, we systematically explored the dosage-dependent impact of cathodal tDCS over the M1, with different stimulation intensities (1, 2, and 3 mA, electrodes size 35 cm^2^), and durations (15, 20, and 30 min), as explored by TMS- elicited motor evoked potentials (MEPs). Low and high-intensity protocols resulted in MEP amplitude reductions, whereas an excitability enhancement was observed after medium intensity tDCS (Mosayebi Samani et al., 2019).

In the present study, we aimed to explore the transferability of these results from the M1 to the PFC. In eight randomized sessions, four cathodal tDCS dosages of low, medium, and high intensity, as well as sham stimulation were applied over the M1, and PFC, with current densities at the scalp-electrode-interface identical to our foregoing study (Mosayebi Samani et al., 2019). tDCS after-effects were then evaluated using TMS-EEG, and TMS-MEP approaches at the regional and global scalp level for TEP and MEP amplitude changes, and TMS-evoked oscillations. In line with recent findings, we further assessed the association between cortical and corticospinal excitability alterations (Ahn and Fröhlich, 2021), as well as tDCS- induced electrical fields (EFs) (Laakso et al., 2019; Mosayebi-Samani et al., 2021). Based on previous findings, we anticipated that all active protocols would modulate TEP amplitudes compared to the respective baseline and/or sham conditions. We also anticipated dosage-dependent nonlinear TEP and MEP amplitude modulations for motor cortex tDCS. Furthermore, we expected gradually different patterns of the neurophysiological effects of tDCS over the PFC, as compared to M1 tDCS.

## 2. Material and Methods

### 2.1. Participants

Since this is the first study investigating the transferability of tDCS-generated cortical excitability alterations from motor to prefrontal cortices, a literature-based, a priori sample size estimation could not be executed. Thus, a post hoc power calculation was performed using G*Power 3.1 (Faul et al., 2009). The analysis was based on a repeated measures ANOVA with a medium to large effect size ƒ = 0.35 (corresponding to the average empirically obtained effect size

**η_p_^2^** = 0.11, see table 2), α = .05, and a sample size of 18 participants (11 males and 7 females, mean age 26.61±3.56 years). This resulted in a power of 0.96. All participants were young, healthy, and right-handed according to the Edinburgh handedness inventory (Oldfield, 1971) and had no history of neurological and psychiatric diseases, or fulfilled exclusion criteria for noninvasive electrical or magnetic brain stimulation (Rossi et al., 2009; Bikson et al., 2016). The study conformed to the Declaration of Helsinki and was approved by the local Ethics Committee of the Leibniz Research Centre for Working Environment and Human Factors. All participants gave written informed consent before starting the study and were financially compensated for participation.

### 2.2. Navigated TMS-EEG and -MEP Measures

#### 2.2.1. Transcranial Magnetic Stimulation

Single-biphasic TMS pulses were applied at 0.33 Hz (±30% jitter) delivered by a PowerMag magnetic stimulator (Mag&More, Munich, Germany) with a figure-of-eight coil (PMD70) held tangentially over the EEG cap, with the handle pointing backwards and laterally at 45° from the midline. For the M1 stimulation site, TMS pulses were first applied to determine the representational area of the right abductor digiti minimi (ADM) muscle, in which the largest MEPs were produced by a given medium TMS intensity. At this stimulation site, the resting motor threshold (RMT) was then determined by the TMS-Motor-Threshold-Assessment Tool (MTAT 2.0) (Awiszus and Borckardt, 2012). For the PFC stimulation site, the TMS coil was placed over the F3 position (10-10 international EEG standard), with the handle pointing backwards and laterally at 45° from the midline. Note that the F3 positioning was selected to approximate the scalp location overlaying the left DLPFC, in accordance with previous studies (Herwig et al., 2003; Beam et al., 2009; Hill et al., 2017).

At each stimulation site, and for each time point, 120 single TMS pulses were applied with a stimulation intensity of 100% RMT. RMT was obtained for the M1 stimulation site with the TMS coil placed at about 5 mm distance to the surface of the head, due to the thin layer of Ten-20 paste, the tDCS electrode and the EEG cap between the coil and the skin. For the PFC stimulation site, we first measured RMT over the M1, with the TMS coil attached to the EEG cap and with a 4 mm thick foam in between, to keep the coil-to-head distance similar to that of the PFC stimulation site with a coil-to-head distance of about 5 mm because of the Ten-20 paste, the tDCS electrode and the EEG cap between the coil and the skin. The foam was then removed and the TMS coil was placed over the PFC stimulation target (F3). TMS intensity was 100% of RMT, to obtain reasonable effects of TMS on EEG responses (Lioumis et al., 2009), but minimize non-direct/unwanted effects of TMS, such as TMS-related artifacts (Mutanen et al., 2013) (decay and muscles artifacts; see section 2.4.1. ‘TEP preprocessing’), and the TMS-induced clicking sound and coil vibration, which result in contamination of the EEG response with sensory and somatosensory responses (Conde et al., 2019; Rocchi et al., 2021).

#### 2.2.2. EEG Recording

A TMS-compatible EEG system (NeurOne, BittiumCorporation, Finland) was used to continuously record TEPs (DC-1.25kHz) with a sampling frequency of 5 kHz. EEG signals were captured by TMS-compatible Ag/AgCl C-ring electrodes via a 64 electrode EEG cap (EasyCap, Herrsching, Germany). Electrodes were positioned according to the 10-10 international EEG standard. For both tDCS stimulation sites, eight EEG electrodes were excluded from data analysis due to the placement of the tDCS electrodes, including: C1, C3, C5, FC3, CP3, Fp2, AF4, and AF8 (for tDCS over M1), and F1, F3, F5, FC3, AF3, Fp2, AF4, and AF8 (for stimulation over the PFC)). Electrodes were online referenced and grounded to external electrodes placed on the forehead (above the nasion). Two additional electrodes were used to record horizontal and vertical eye movements (one on the orbital ridge centered directly below the left eye and the other one at the lateral junction of the upper and lower right eyelids) (Fecchio et al., 2017). Impedances of all electrodes were kept below 5 kΩ during the experiment.

#### 2.2.3. Navigated TMS

Following individual TMS stimulation site identification (see section 2.2.1), an MR-based 3D-navigation system (PowerMAG View, Mag&More, Munich, Germany) was employed to store and display online the position and orientation of the TMS coil with respect to the participant’s head and fiducials based on an individual structural MRI scan, assuring accuracy and reproducibility of the stimulation outcome throughout the experiment (Tremblay et al., 2019). All imaging data were acquired by a 3T Philips Achieva scanner (Best, Netherlands) with a 32-channel head coil. Anatomical images were recorded based on T1-weighted fast 3D gradient echo pulse sequences (repetition-time= 8179 ms, echo-time= 3.7 ms, flip-angle= 8°, 220 slices, matrix-size= 240x240, and resolution= 1x1x1 mm^3^).

#### 2.2.4. MEP Recording

Surface EMG was recorded from the right ADM in a belly-tendon montage. The signals were amplified (Gain: 1000), band-pass filtered (2 Hz–2 kHz) using D440-2 (Digitimer, WelwynGardenCity, UK), and digitized (sampling rate: 5 kHz) with a micro 1401 AD converter (CED, Cambridge, UK), controlled by Signal Software CED, v.2.13.

#### 2.2.5. Transcranial Direct Current Stimulation

tDCS was applied with a constant-current battery-powered stimulator (neuroCare, Ilmenau, Germany), through a pair of surface rubber electrodes (25 cm^2^) attached on the scalp using conductive paste (Ten20^®^, Weaver). For the M1 stimulation sessions, the target electrode was centered over C3 and rotated 45^0^ towards the midline. For the PFC stimulation sessions, the target electrode was centered over the F3 position, parallel to the midline. The return electrode, for both stimulation sites, was located on the contralateral supraorbital region. Prior to electrode placement, a topical anesthetic cream (EMLA^®^, AstraZeneca, UK) was applied to the respective stimulation sites, in order to decrease somatosensory sensations and sufficiently blind the participants (McFadden et al., 2011). In eight randomized sessions, four cathodal tDCS dosages of low (0.7 mA for 15 min), medium (1.4 mA for 20 min), and high (2.1 mA for 20 min) intensities, and sham (for 15 min), were applied over the two stimulation sites M1 and PFC. We chose stimulation parameters (intensity and duration) based on the results of our former study (Mosayebi Samani et al., 2019), but used smaller stimulation electrodes (25 cm^2^ instead of 35 cm^2^) to sacrifice fewer EEG channels around the tDCS target sites. Therefore, the same current densities at the scalp-electrode interface as in the previous study were applied for low (0.028 mA/cm^2^), medium (0.057 mA/cm^2^), and high (0.085 mA/cm^2^) dosages of tDCS. For sham stimulation, 0.7 mA stimulation was delivered for 15 s followed by 15 min with 0.0 mA stimulation.

All protocols were conducted with a 10 s ramp-up and -down at the start, and end of stimulation. The tDCS sessions were applied in randomized order with a minimum of seven days between sessions to avoid carry-over effects (Nitsche et al., 2008).

### 2.3. Experimental procedures

The study was performed in a cross-over single-blinded sham-controlled repeated measures design. At the beginning of each session, participants were seated in a comfortable chair with head- and arm-rests. Afterwards, the topical anesthetic cream was applied over the scalp at the identified corresponding tDCS stimulation sites, and the tDCS electrodes were attached to the head with conductive paste (see section 2.2.5 for details), followed by the set-up of the EEG cap. The participant’s head was then co-registered to the individual head model within the navigation system. Thereafter, TMS stimulation sites were identified for M1 and PFC stimulation and stored in the navigation system, to be used throughout the experiment (see section 2.2.1 for details). Subsequently, TMS intensity was adjusted to the RMT (as explained above). Then baseline cortical excitability was determined by TMS-EEG over the M1 (including also simultaneous MEP recording) or PFC stimulation sites, depending on the tDCS stimulation session. Afterwards, the respective tDCS protocol was applied. Finally, cortical excitability was monitored by TMS-EEG immediately (POST0), 30 min (POST30), 60 min (POST60), and 120 min (POST120) after tDCS, Figure 1. Concurrent with TEP recording, the participants were exposed to white noise through headphones to minimize contamination of the EEG signal by auditory evoked potentials, induced by the TMS-related clicking sound (Nikouline et al., 1999; ter Braack et al., 2015; Rocchi et al., 2021). Here, prior to the start of the main experiment, the loudness of the ‘noise masking’ sound was systematically measured (using Sound level Analyzer B&K 2250, Brüel & Kjær, Denmark) and respective values were then used for controlling the upper safety threshold (95db). During the experiment, the sound level was gradually increased until the participant could not hear the “click” sound of the TMS coil, or until it had reached the upper safety threshold (Rocchi et al., 2021).

**Figure 1.**
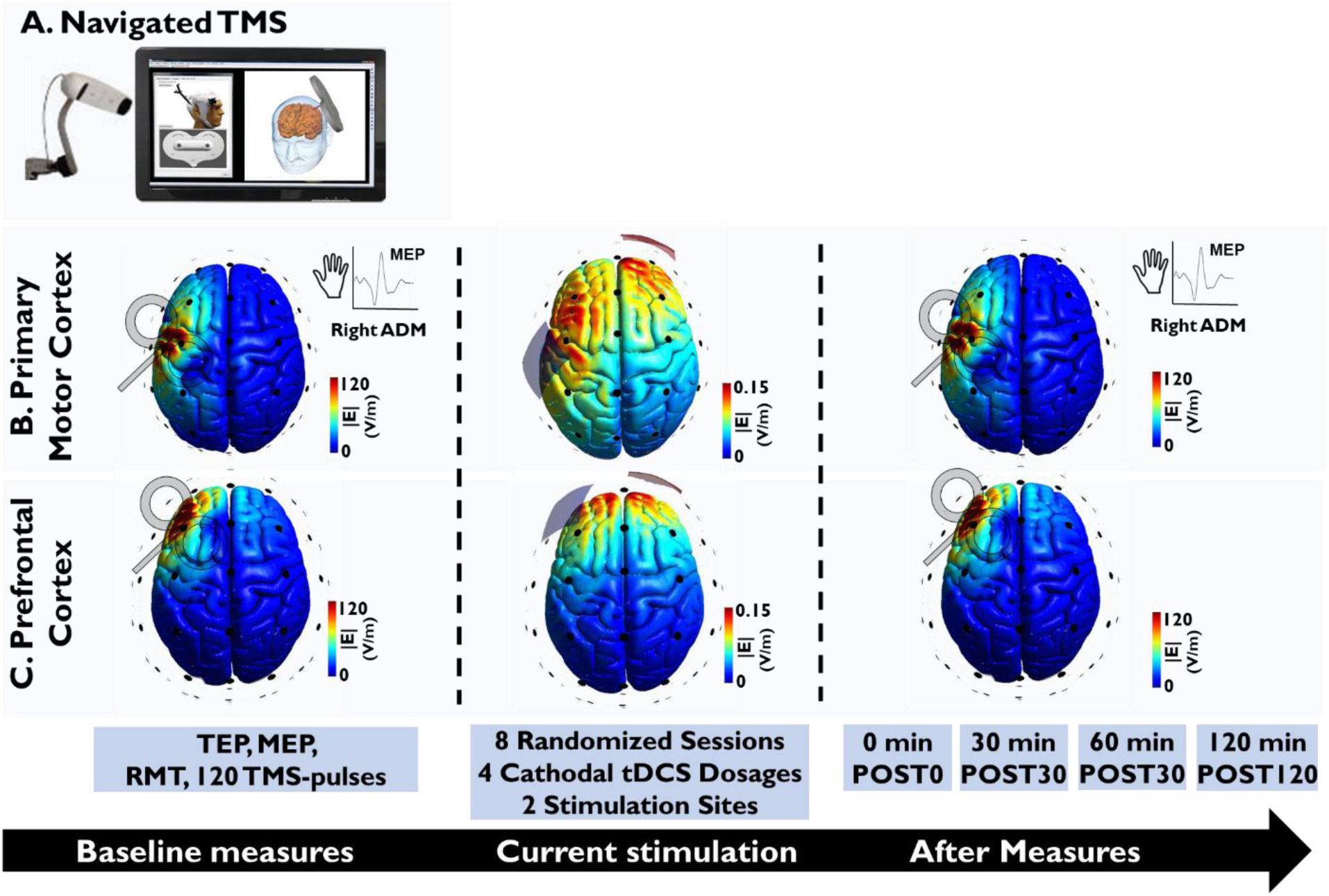
**Course of the study**. In eight randomized sessions, four cathodal tDCS dosages of low (0.028 mA/cm2; 15 min), medium (0.057 mA/cm2; 20 min), and high (0.085 mA/cm2; 20 min) intensity, as well as sham stimulation (0mA; 15min) were applied to target the primary motor (M1; **B**), and dorsolateral prefrontal cortex (PFC; **C**). To evaluate the modulatory effects of tDCS, using a navigated TMS system (**A**), TMS-evoked cortical reactivity, and TMS-elicited MEP (only for tDCS applied over the M1), were recorded before tDCS, and immediately (POST0), 30 min (POST30), 60 min (POST60), and 120 min (POST120) after tDCS, with 120 TMS pulses at each time-point. The recorded data were then evaluated for TMS-evoked potentials (TEPs), oscillations, and MEP alterations. Colors in the cortical grey matter are illustrating electric field magnitudes induced by TMS and tDCS (0.7 mA) estimated via SimNIBS open-source software with its default parameters and head model (erni.msh) (Thielscher et al., 2015).

### 2.4. Calculations

Offline data processing was performed with custom scripts in MATLAB (R2019b, Mathworks, USA), Fieldtrip (Oostenveld et al., 2011), Brainstorm (Tadel et al., 2011), and FreeSurfer (Dale et al., 1999) toolboxes. All custom codes are available at Extended Data 1.

#### 2.4.1. MEP and TEP preprocessing

***MEP*** amplitudes were first visually inspected to exclude MEP trials: 1) in which background electromyographic activity was present, and 2) which were associated with respective bad TEP trials (see below). Then, the individual means of peak-to- peak MEP amplitudes, recorded at each time-point, were separately calculated for all subjects and all conditions.

***TEP*** *preprocessing*: first, a time period (-5 to +15ms) around each TMS pulse was removed and interpolated. Then the EEG data were segmented into epochs around the TMS pulses (−1000 to 1500 ms), visually inspected to remove bad trials/channels (Ilmoniemi and Kicic, 2010), and referenced to the average of all electrodes. Afterwards, the data were high-pass filtered (1Hz; 4th-order zero-phase Butterworth) and preprocessed with the ‘signal-space projection with source-informed reconstruction’ (SSP-SIR) algorithm. SSP-SIR is a spatial filtering method, which has been shown to efficiently suppress TMS-related muscle artifacts (Mutanen et al., 2013; Mutanen et al., 2016). For that, we formed subject-specific, realistic lead-field matrices, by first automatically segmenting the individual T1- weighted MRI images using Freesurfer software (Dale et al., 1999), which were then imported to Brainstorm toolboxes (Tadel et al., 2011), to generate lead-fields, based on the three-layer symmetric boundary element method via OpenMEEG (Gramfort et al., 2010) (tissue conductivity values (S/m): scalp = 0.33, skull = 0.0033, and brain = 0.33; standard 10-10 EEG electrode location adapted by MR-based participant’s fiducials). Then, SSP-SIR was applied to project out artifacts during the first 50 ms after the TMS pulse. We identified the components that explained more than 90% of the variance of the high-pass filtered data (above 100Hz) in the EEG trials and removed those from the TEP data (Mutanen et al., 2016). Data were then low-pass filtered (100Hz; 4th-order zero-phase Butterworth) following an independent component analysis (FastICA) to remove remaining noise components (Hyvärinen and Oja, 2000; Hernandez-Pavon et al., 2012; Rogasch et al., 2014; Rogasch et al., 2017). Finally, the decomposed data were filtered (lowpass: 45 Hz; 4th-order, zero-phase Butterworth), baseline-corrected (baseline EEG was obtained −1000 to −50 ms relative to TMS onset; a period not closer than -50ms to the TMS pulse was chosen to avoid contamination from the TMS artifact (Gordon et al., 2018b)), and bad channels were interpolated (according to the distance from neighboring channels; note that only bad EEG channels were interpolated, but not the EEG channels that were removed due to the placement of tDCS electrodes; see section 2.2.2). For the included number of trials, removed MEPs, removed ICA components, and interpolated EEG channels see Table 1-1.

**Table 1.**
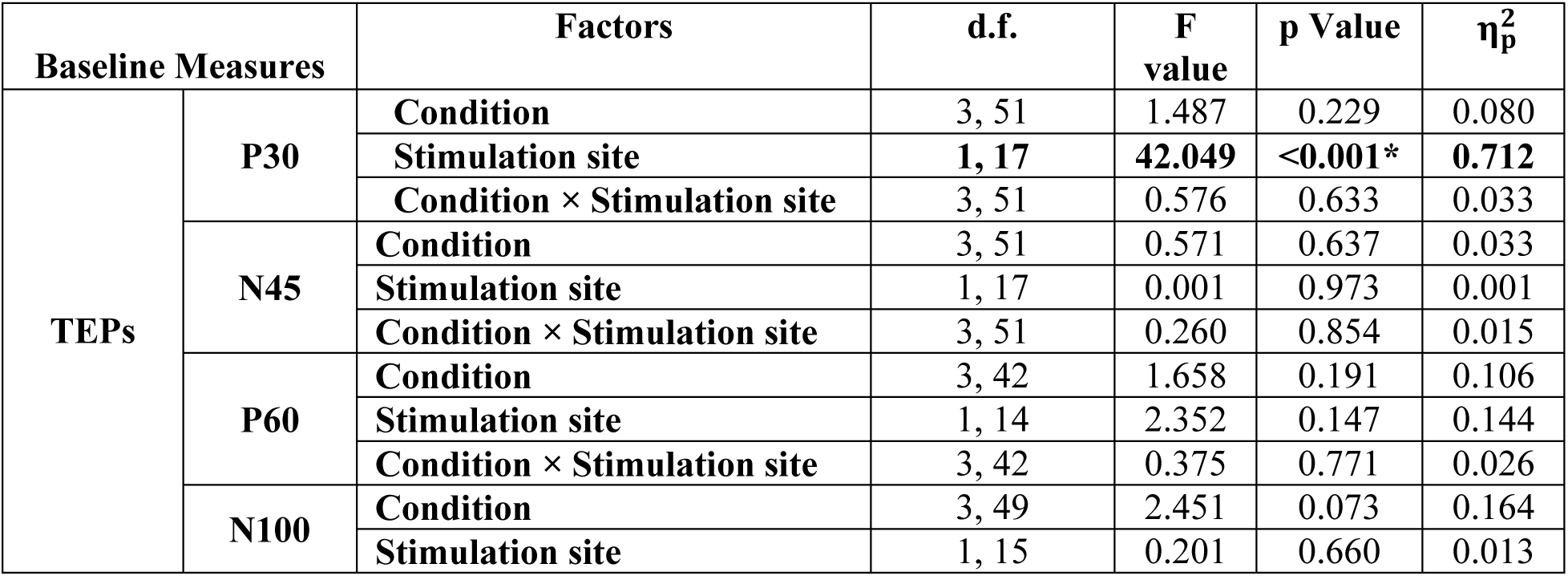
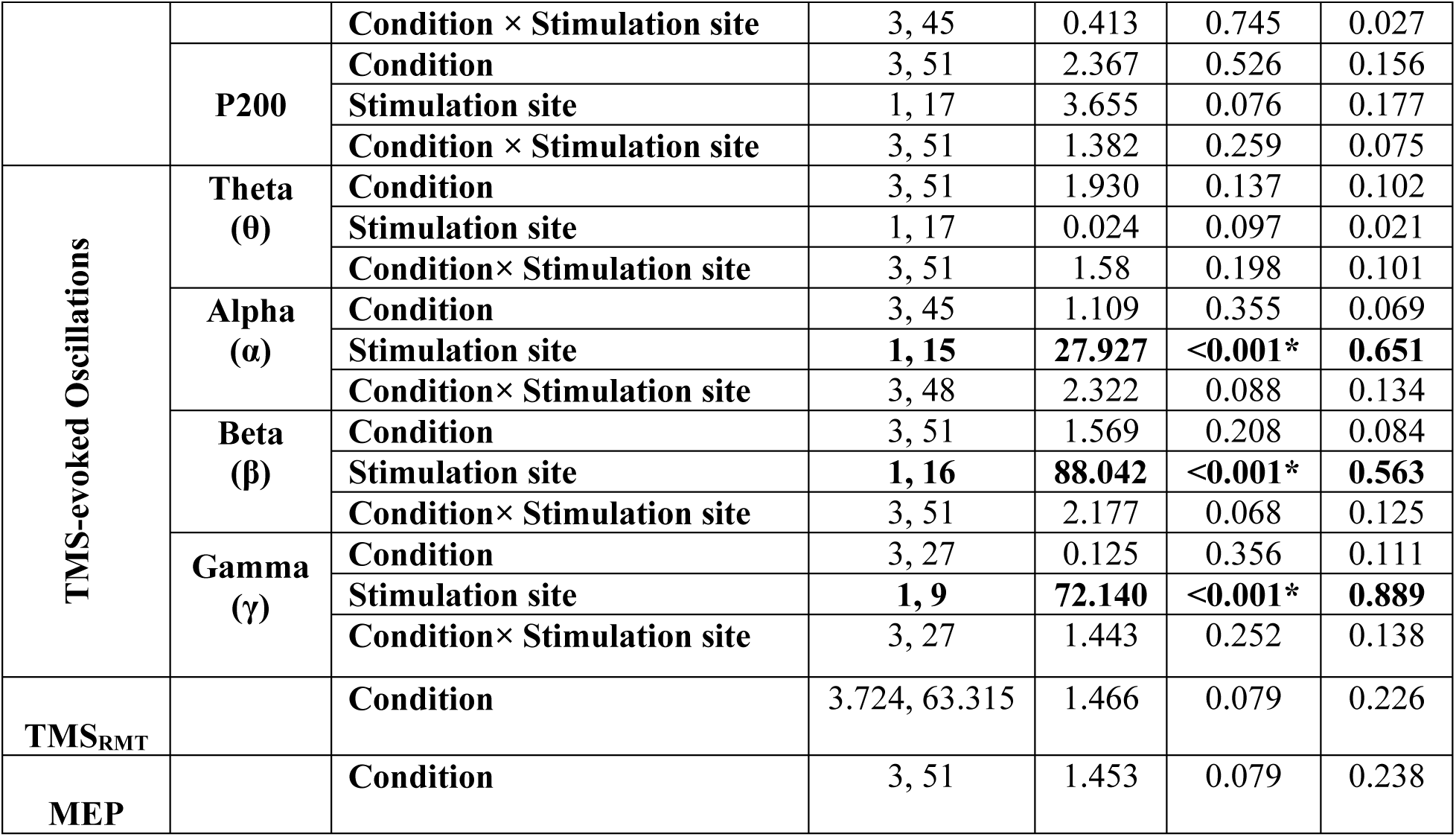
Results of the ANOVAs conducted to evaluate baseline measurements. The ANOVAs showed no significant differences of baseline measures within stimulation sites; however, significant differences were observed for the TEP P30 between stimulation sites, and FB for all frequency bands, except theta. Asterisks indicate significant effects (p < 0.05), d.f. = degrees of freedom, η*p*^2^ = partial eta squared, TEP = TMS-evoked Potentials and FB = frequency band. See also Table 1-1 for the TEPs and MEPs included in the data.

#### 2.4.2. TMS-evoked Potentials

To evaluate the regional effects of tDCS applied over M1, and the PFC, we averaged the TEP deflections measured by the FC1 and CP1 electrodes (region of interest (*ROI*_*M*1_), and the FCz and Fz electrodes (*ROI*_*PFC*_ ), respectively. These ROIs were selected to capture the regional effects of tDCS, as they are located close to the tDCS target electrode, are distant from cranial muscles, which are a source of TMS-related artifacts (Mutanen et al., 2013; Hill et al., 2017), and do not overlap between the two tDCS stimulation sites (M1 and PFC).

For these ROIs, the known TEP peaks were first identified by searching the maximum (for positive) or minimum value (for negative deflections). TEP deflections were identified for the following time periods of 20-40 ms (P30), 35-55 ms (N45), 45-75 ms (P60), 85-135 ms (N100) and 170-230 ms (P200) (Komssi and Kahkonen, 2006; Lioumis et al., 2009; Rogasch and Fitzgerald, 2013; Tremblay et al., 2019). A 10 ms window (±5 ms) around each identified TEP peak was then averaged to calculate the respective TEP amplitude, to be used for further statistical analysis (Hill et al., 2017).

In addition, respective analyses were also performed to explore *global effects* of tDCS, within all available EEG electrodes across all individuals using a global scalp analysis (see statistical methods (2.8.2.2) for details).

#### 2.4.3. TMS-evoked Oscillations

To test if tDCS modulated TMS-related neural oscillations, time-frequency representations (TFRs) of oscillatory power were calculated for *ROI*_*M*1_ and *ROI*_*PFC*_ on a single trial basis (Morlet wavelet; wavelet width: starting from 2.6 cycles and adding 0.2 cycle for each 1 Hz), and then normalized (db) to the respective baseline (−500 to −100 ms). Finally, frequency power estimates were calculated for four separate frequency bands (FBs), including Theta (θ; 4-7 Hz), Alpha (α; 8-13 Hz), Beta (β; 14-29 Hz) and Gamma (γ; 30-45 Hz) within the time window of 50–300 ms after the TMS pulse for all frequency bands (Gordon et al., 2018b).

In addition, respective analyses were also performed to explore *global effects* of tDCS, within all available EEG electrodes across all individuals using a global scalp analysis (see statistical methods (2.8.2.4) for details).

#### 2.4.4. Discriminability and qualitative assessment of tDCS protocols

After finishing each session, participants were asked to fill in a questionnaire which contained: 1. Guessed intensity of tDCS (none, low, medium, high) 2. Rating scales for evaluation of the presence and amount of visual phenomena, itching, tingling, and pain during stimulation, and 3. Rating scales for evaluation of the presence and amount of skin redness, headache, fatigue, concentration difficulties, nervousness, and sleep problems within 24 hours after stimulation. The side-effects were rated on a numerical scale ranging from zero to five, zero representing no and five extremely strong sensations (Poreisz et al., 2007; Brunoni et al., 2011).

#### 2.4.5. Computational modeling of tDCS- and TMS-induced Electrical Fields

Using the individual T1 image of each participant, the tDCS- and TMS-induced EFs were estimated with a free and open source software package for simulation of non-invasive brain stimulation (SimNIBS v.3.2.3) (Thielscher et al., 2015). Briefly, this includes T1 image segmentation into the major head tissues, 3D volume reconstruction, placement of tDCS electrodes (for tDCS-induced EF estimation: C3- Fp2 for M1 stimulation and F3-Fp2 for tDCS over the PFC; electrode size 5*5 cm^2^), or locating the TMS coil (figure-of-8 shape, 5mm above the head; to approximate the actual experimental condition; 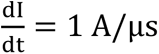), assigning the respective tissue conductivities (white matter: 0.15 S/m, gray matter (GM): 0.4 S/m, CSF: 1.79 S/m, eyeballs and scalp: 0.33 S/m, skull: 0.008 S/m), and calculating the tDCS- (for low (0.7 mA), medium (1.4 mA), and high-dosage (2.1 mA) intensity)- and TMS-induced EFs by means of the finite element method, under the quasi-static approximation (Miranda et al., 2013). Then, for each participant and for each stimulation site (M1 and PFC), a mask was created over the GM, where the TMS-induced EF was 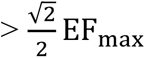 (the half power region, which is a standard measure of the focality of TMS-induced EFs over the targeted area (Carbunaru and Durand, 2001; Deng et al., 2013)). Finally, the tDCS-induced EFs (strength: |E|; 95% percentile) over the masks were calculated (Figure 11.C, Figure 11-1, Figure 11-2).

**Figure 2.**
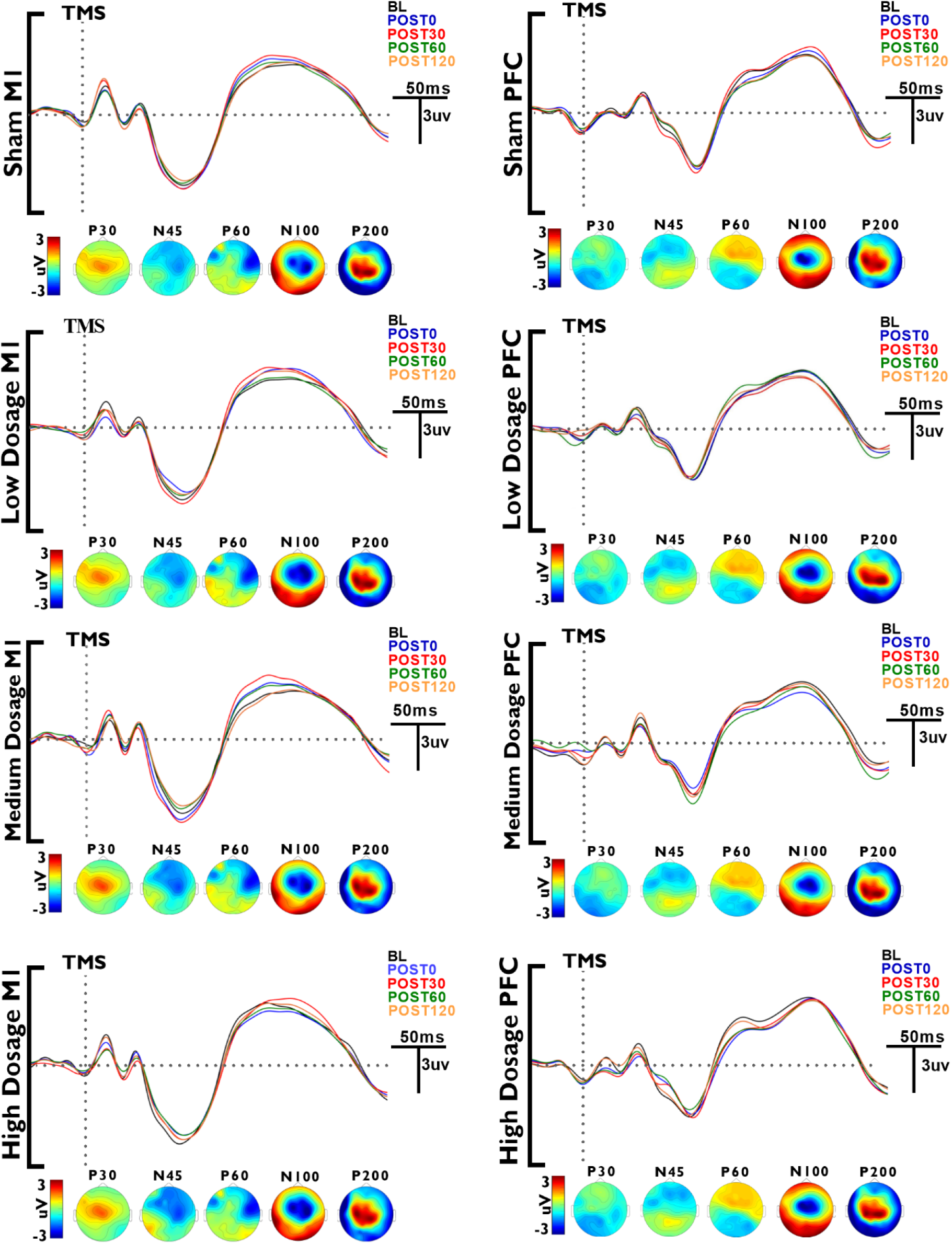
Local effects of tDCS on TMS-evoked Potentials. Cathodal tDCS dosages of low, medium, and high intensities, and sham, were applied over the primary motor (M1) and left dorsolateral prefrontal cortex (PFC) stimulation sites. Local tDCS effects were obtained immediately (POST0) to up to two hours after stimulation (POST30, POST60, POST120), over the ROI_M1_ (averaged FC1 and CP1 electrodes) and ROI_PFC_ (averaged FCz and Fz electrodes). Grand average across all subjects following tDCS conditions over the left primary motor cortex (ROI_M1_; left column) and left DLPFC (ROI_PFC_; right column), and topographic plots displaying voltage distributions across the scalp for each TEP peak ((P30, N45, P60, N100, P200) at the respective stimulation sites are shown.

### 2.8. Statistics

All statistical analyses were performed using SPSS (IBM Corp. v.26.0), custom scripts in MATLAB, and the Fieldtrip toolbox (Oostenveld et al., 2011).

#### 2.8.1. Baseline measures

At the regional level (*ROI*_*M*1_, *ROI*_*PFC*_ ), to test if baseline measures differed between sessions within and/or between stimulation sites, two-way repeated-measures ANOVAs (rmANOVA) were performed, with condition (4 levels) and stimulation site (2 levels) as within-subject factors, and baseline of TEP peak, or TFR of each frequency band, as dependent variables. In addition, two separate one-way rmANOVAs were performed with condition (8 levels for TMS_RMT_, 4 levels for baseline MEP) as within-subject factor, and TMS_RMT_ intensity, or baseline MEP as dependent variables, respectively.

Furthermore, at the global scalp level, to test if baseline measures were comparable between tDCS sessions within stimulation sites, cluster-based nonparametric permutation tests were used, based on the Monte-Carlo method (Maris and Oostenveld, 2007), to effectively control for multiple comparisons across numerous EEG channels and time-points (Maris and Oostenveld, 2007). The clusters were defined as ≥2 neighboring electrodes with a p-value<0.05 (t-test), the number of permutations was 5000, and Monte-Carlo two-tailed p-values were defined for a critical α level of p<0.05 (Hill et al., 2017; Rogasch et al., 2020). For the TMS-evoked potentials we selected a time period of 20-250ms after TMS and for TMS-evoked oscillatory data a time period of 50–300ms for all frequency bands (Gordon et al., 2018b). These time intervals were selected to avoid any *a priori* assumptions and also improve the methodological compatibility with the regional analysis. For the permutation tests within each stimulation site, we have excluded the missing EEG electrodes due to the placement of tDCS electrodes (8 electrodes for each stimulation site; see section ‘2.2.2. EEG Recording’ for details). However, for comparisons between stimulation sites, this would result in a high total number of missing EEG electrodes (N=13). We therefore conducted the global scalp analysis for each stimulation site separately, with otherwise identical parameters, except for the included channels. The same parameters were used for the further global scalp analysis.

#### 2.8.2. Modulatory effects of tDCS and motor-to-prefrontal transferability

##### 2.8.2.1. tDCS-altered TMS-evoked Potentials (regional effects)

We investigated if, at the regional level (*ROI*_*M*1_, *ROI*_*PFC*_ ), within each stimulation site, the tDCS after-effects differed vs. baseline values. To this end, three-way rmANOVAs were calculated with condition (4 levels; Low, Medium, High dosage, and Sham), time-point (5 levels; baseline, POST0, POST30, POST60, and POST120), and stimulation site (2 levels; M1 and PFC) as within-subject factors, and the absolute values of each TEP peak (P30, N45, P60, N100, P200) as the dependent variable.

In addition, we investigated if the tDCS-induced after-effects differed from respective sham conditions, between active stimulation conditions within each stimulation site, and effects of the tDCS protocols differed between the M1 and PFC stimulation sites. To this end, individual means of the TEP peaks after tDCS were first normalized (Δ) to the respective individual mean baseline: (*post.tDCS − baseline*) ⁄ |*baseline*|. Then, three-way rmANOVAs were calculated with condition (4 levels), time-point (4 levels; POST0, POST30, POST60, and POST120) and stimulation site (2 levels) as within-subject factors, and the normalized value of each TEP peak (ΔP30, ΔN45, ΔP60, ΔN100, ΔP200), as the dependent variable.

##### 2.8.2.2. tDCS-altered TMS-evoked Potentials (global scalp effects)

To assess the distributed effects of tDCS, we performed a global scalp analysis of TMS-evoked potentials using cluster-based nonparametric permutation tests, based on the Monte- Carlo method (Maris and Oostenveld, 2007); see section ‘2.8.1. Baseline measures’ for details of statistical parameters). We then investigated if within each stimulation site 1) the active intervention conditions altered cortical outcome measures versus baseline and 2) the after-effects of tDCS differed from the sham condition.

##### 2.8.2.3. tDCS-altered TMS-evoked Oscillations (regional effects)

to assess the regional effects of tDCS on TMS-evoked oscillations, the same statistics (rmANOVAs; see section ‘TMS-evoked Potentials (regional effects’)) were performed, but here with TFRs of each FB (Theta, Alpha, Beta, and Gamma) as dependent variables.

##### 2.8.2.4. tDCS-altered TMS-evoked Oscillations (global scalp effects)

To investigate the global effects of tDCS on TMS-evoked oscillations, the same statistics (cluster-based permutation test, see section ‘TMS-evoked Potentials (global scalp effects’) were performed, but here with TFRs of each frequency band (Theta, Alpha, Beta, and Gamma).

##### 2.8.2.5. tDCS-altered TMS-elicited MEPs

we investigated if the tDCS after- effects on MEP amplitudes differed vs. baseline values. To this end, two-way rmANOVAs were calculated with condition (4 levels; Low, Medium, High dosage, and Sham) and time-point (5 levels; baseline, POST0, POST30, POST60, and POST120) as within-subject factors, and absolute values of the MEP measures as the dependent variable. Another two-way rmANOVA with condition (4 levels), and time point (4 levels; POST0, POST30, POST60, and POST120) as within-subject factors, and ΔMEP as the dependent variable, was calculated to test if MEP amplitudes differed vs. sham, and between active stimulation conditions.

##### 2.8.2.6. Association between tDCS-generated TEP or MEP alterations, and tDCS-induced Electrical Fields

To explore associations between tDCS-generated TEP or MEP alterations, and tDCS-induced EFs, we calculated Pearson correlations for these variables. Furthermore, we explored the comparability of tDCS-induced EFs between the two stimulation sites via *Student’s* paired t-tests. Due to the exploratory nature of these analyses, we did not correct for multiple comparisons.

#### 2.8.3. Qualitative assessment of tDCS protocols

To identify if participants correctly guessed tDCS intensities correctly, chi-square tests were conducted. Side-effects during and after tDCS were analyzed by a repeated measure ANOVA with condition (8 levels) as within-subject factor and rating scores (0-5) as the dependent variable. In case of significant effects, follow-up exploratory post-hoc paired t-tests were conducted to examine if an active session resulted in a significantly different sensation relative to sham.

For all ANOVAs, Mauchly’s test of sphericity was conducted, and the Greenhouse-Geisser correction was applied when necessary. The critical significance level was set at *p* ≤ 0.05. Post hoc tests were conducted using the Fisher Least Significant Difference test, in case of significant ANOVA results.

## 3. Results

### 3.1. Baseline measures

The respective rmANOVAs, at the regional level, showed no significant differences of baseline TEP peaks within each stimulation site, however, significant differences between stimulation sites were observed for P30. Post-hoc tests indicated a lower amplitude of the P30 over the PFC, as compared to M1 (Table 1, Figure 2, Figure 3). Also, the respective rmANOVAs showed no significant differences for baseline TFRs of each frequency bands within each stimulation site. However, significant differences were identified between stimulation sites for all frequency bands (except Theta), with lower power over the PFC, as compared to M1 (Table 1, Figure 6, Figure 7). No significant differences were observed for either TMS_RMT_ (Table 1), or baseline MEP amplitudes between stimulation conditions (Table 1, Figure 10).

**Figure 3.**
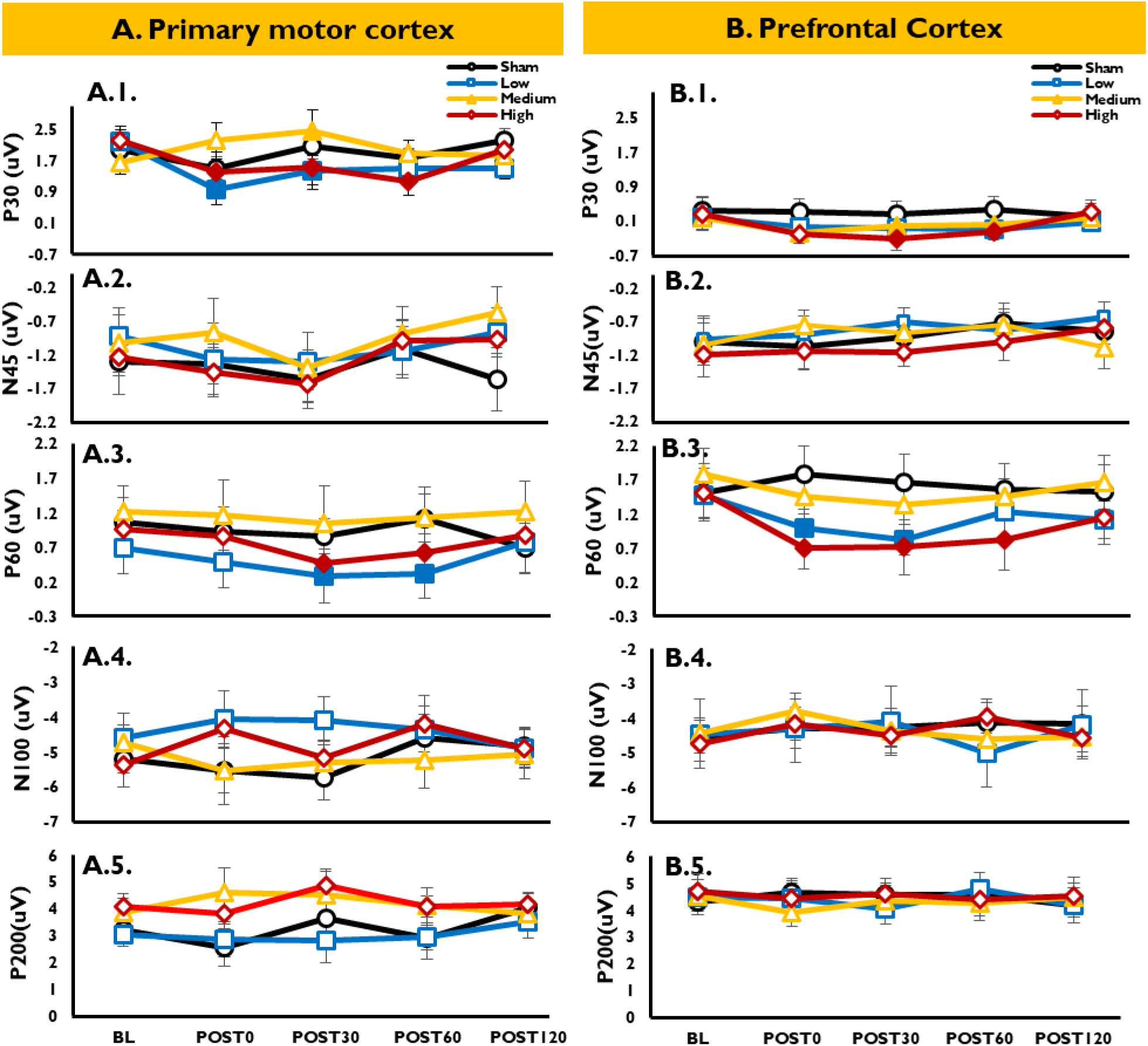
Local effects of tDCS on TMS-evoked Potentials (absolute values; comparison vs. baseline). Cathodal tDCS dosages of low, medium, and high intensities, and sham, were applied over the primary motor (M1) and left dorsolateral prefrontal cortex (PFC). The local tDCS effects were then evaluated immediately (POST0) for up to two hours after stimulation (POST30, POST60, POST120), over the ROI_M1_ (averaged FC1 and CP1 electrodes) and ROI_PFC_ (averaged FCz and Fz electrodes). **A1-5, B1-5:** absolute TMS-evoked potentials over the M1 and the PFC, respectively, are shown. tDCS generated a dosage- dependent, partially non-linear modulation of TEP (P30, N45, P60, N100, P200) over the different stimulation sites, as shown by the amplitude alterations of early (P30 and P60) TEP peaks. Filled symbols show a significant difference of active tDCS conditions vs. baseline. Error bars show the standard error of the mean (SEM). See figure 3-1 for the local effects of tDCS on TMS-evoked Potentials (normalized values).

Also, at the global scalp level, using cluster-based permutation tests, no significant differences were observed between baseline measures of different tDCS dosages within each stimulation site.

### 3.2. Effects of tDCS and motor-to-prefrontal transferability

#### 3.2.1. tDCS-altered TMS-evoked Potentials (regional Effects)

For the P30 TEP, the primary rmANOVA (absolute values), conducted to explore the modulatory effects of tDCS, resulted in significant main effects of time-point and stimulation site, interactions of ‘condition × time-point’, ‘time-point × stimulation site’ and ‘condition × time-point × stimulation site’ (F_(5.260,79.012)_ = 2.311, p = 0.045, ɳ^2^ = 0.134), but no significant main effect of condition, or other interactions (Table 2 and Figure 2, Figure 3). For **M1 stimulation,** post-hoc tests comparing tDCS after-effects with baseline showed a significant P30 amplitude reduction for low- (POST0, POST30) and high-dosage stimulation (POST0, POST30, POST60), while medium-dosage tDCS increased the TEP amplitude (only at POST30) for M1. For **PFC stimulation,** the post hoc tests showed a significant P30 amplitude reduction for medium-(POST0, POST30, and POST60) and high-dosage (POST30, and POST60) tDCS.

**Table 2.**
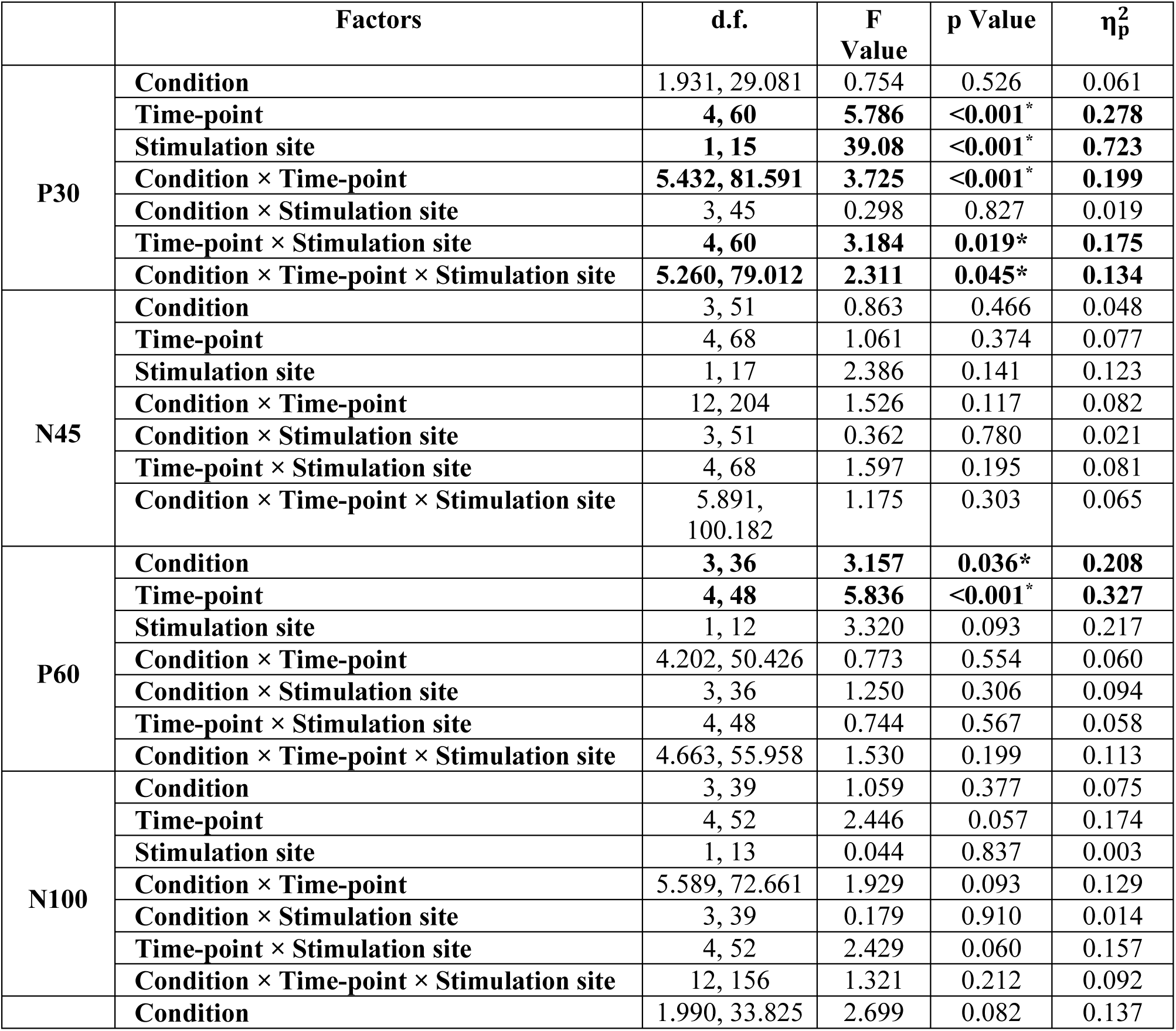
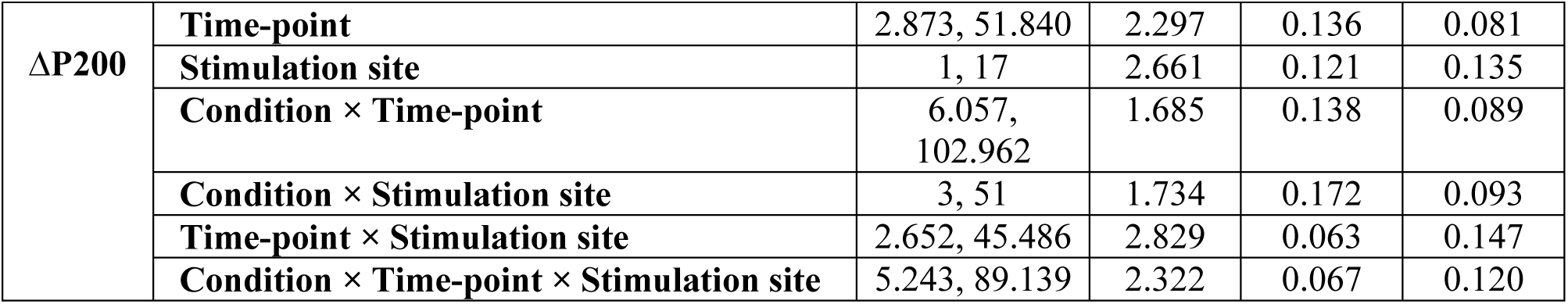
Results of the ANOVAs conducted for tDCS-induced TEP alterations (absolute values). The statistical results indicate tDCS-induced effects for the early (P30 and P60) TEP peaks, with no one-to-one transferability of tDCS effects from the motor to the prefrontal cortex. Asterisks indicate significant effects (p < .05), d.f. = degrees of freedom, η*p*^2^ = partial eta squared. See also Table 2-1 for the results of the ANOVAs conducted for tDCS-induced TEP alterations (normalized values).

In addition, the secondary rmANOVA conducted with standardized values to compare the effects of each tDCS condition with sham, between active stimulation conditions within each stimulation site, and differences of tDCS effects between the M1 and PFC stimulation sites, showed significant main effects of condition and time-point and a significant interaction of ‘condition × time-point’, but no main effect of stimulation site, or significances of other interactions (Table 2-1, Figure 3- 1). Post-hoc tests comparing tDCS after-effects at the respective time points vs sham showed a significant ΔP30 amplitude reduction for low- (POST0) and high-dosage stimulation (POST0, POST30, POST60), while medium-dosage tDCS increased the TEP amplitude (POST0, POST30) for M1 tDCS. For **PFC stimulation**, the results showed a significant ΔP30 amplitude reduction for medium- and high-dosage (both at POST30 and POST60) tDCS, as compared to the respective sham condition (Figure 3-1, Table 2-1). In addition, post-hoc tests comparing active condition, within each stimulation site, showed a larger ΔP30 amplitude for medium-dosage M1 tDCS (POST0, POST30, POST60), as compared to low- and high-dosage M1 tDCS. Finally, post-hoc tests comparing conditions **between stimulation sites** indicated a significant difference for medium-dosage tDCS (POST30, POST60), with a larger P30 amplitude over the motor cortex, as compared to the prefrontal cortex (Figure 3-1).

For the N45 TEP, the primary and secondary rmANOVAs indicated no significant effects of tDCS protocols compared to baseline (Table 2, Figure 2, and Figure 3) and sham values (Figure 3-1, Table 2-1).

For the P60 TEP, the primary rmANOVA resulted in significant main effects of condition and time-point, but no significant main effect of stimulation site, or significances of interactions (Table 2, Figure 2, and Figure 3). Post-hoc tests comparing tDCS after-effects to baseline, **for M1**, indicated a significant P60 amplitude reduction for low and high dosages (both at POST30 and POST60), and **for the PFC** showed a significant P60 amplitude reduction for low- ( POST0, POST30) and high-dosage tDCS (POST0,POST30, and POST60). In addition, the secondary rmANOVA resulted in a significant main effect of condition and a significant interaction of ‘condition × time-point × stimulation site’ (F_(4.648,55.777)_ = 2.185, p = 0.035, ɳ^2^ = 0.122), but no significant effects of the other main factors or interactions (Table 2-1, Figure 3-1). Post-hoc tests comparing tDCS after-effects to sham at the respective time points showed a significant ΔP60 amplitude reduction for low- and high-dosage stimulation over M1 (POST60)**. For PFC stimulation,** the ΔP60 amplitude was significantly reduced vs sham for low- (POST0) and high-dosage tDCS (POST0 and POST30). In addition, post-hoc tests comparing active conditions within each stimulation site showed a larger ΔP60 amplitude for medium-dosage M1 tDCS ( POST60), as compared to low- and high-dosage M1 tDCS; the same post-hoc tests conducted for tDCS over the PFC showed a larger ΔP60 amplitude for medium-dosage PFC tDCS ( POST0), as compared to low- and high-dosage PFC tDCS (Table 2-1, Figure 3-1).

For the N100 and P200 TEP, the respective rmANOVAs showed no significant effects of tDCS protocols comparing to baseline (Table 2, Figure 2, Figure 3) or sham values (Table 2-1, Figure 3-1).

In summary, the results showed non-linear dosage-dependent regional after-effects of tDCS over M1, as indicated by amplitude changes on early TEP peaks (P30 and P60) as compared to baseline, as well as compared to sham measures. However, a rather uniform reduction of early positive peaks was found for the tDCS after-effects over the PFC. No significant effects were however identified for the influence of tDCS on late TEP peaks in M1 or PFC tDCS conditions.

#### 3.2.2. tDCS-altered TMS-evoked Potentials (global scalp effects)

At the global scalp level, cluster-based permutation tests comparing the tDCS after- effects with respective baseline measures showed a negative cluster over the central electrodes (POST0; p=0.024, time-period: 23-48 ms after TMS) and another negative cluster over the parieto-occipital electrodes (POST0; p=0.029, time-period: 170-245 ms after TMS) for low-dosage ***tDCS over M1***). Likewise, a negative cluster was identified over the central electrodes (POST30; p=0.024, time-period: 24-61 ms after TMS) and another negative cluster was revealed over the parieto-occipital electrodes (POST30; p=0.034, time-period: 151-251 ms after TMS). For medium- dosage tDCS, a positive cluster was identified only over the central electrodes (POST30; p=0.041, time-period: 21-54 ms after TMS). In addition, for high-dosage tDCS, a negative cluster was identified over the central electrodes (POST0; p=0.028, time-period: 22-61 ms after TMS), over the parietal electrodes (POST0; p=0.029, time-period: 70-148 ms after TMS), and another negative cluster was identified over the central electrodes (POST30; p=0.024, time-period: 21-60 ms after TMS); Figure 4. Moreover, the results of cluster-based permutation tests comparing the active tDCS conditions with sham measures showed a positive cluster for medium-dosage tDCS over M1 over the central electrodes (POST30; p=0.041, time-period: 30-47 ms after TMS), a negative cluster for high-dosage tDCS over M1 over the centro- medial electrodes (POST0; p=0.032, time-period: 24-65 ms after TMS) and another negative cluster over the centro-medial electrodes (POST30; p=0.021, time-period: 31-86 ms after TMS), Figure 4-1.A-C.

**Figure 4.**
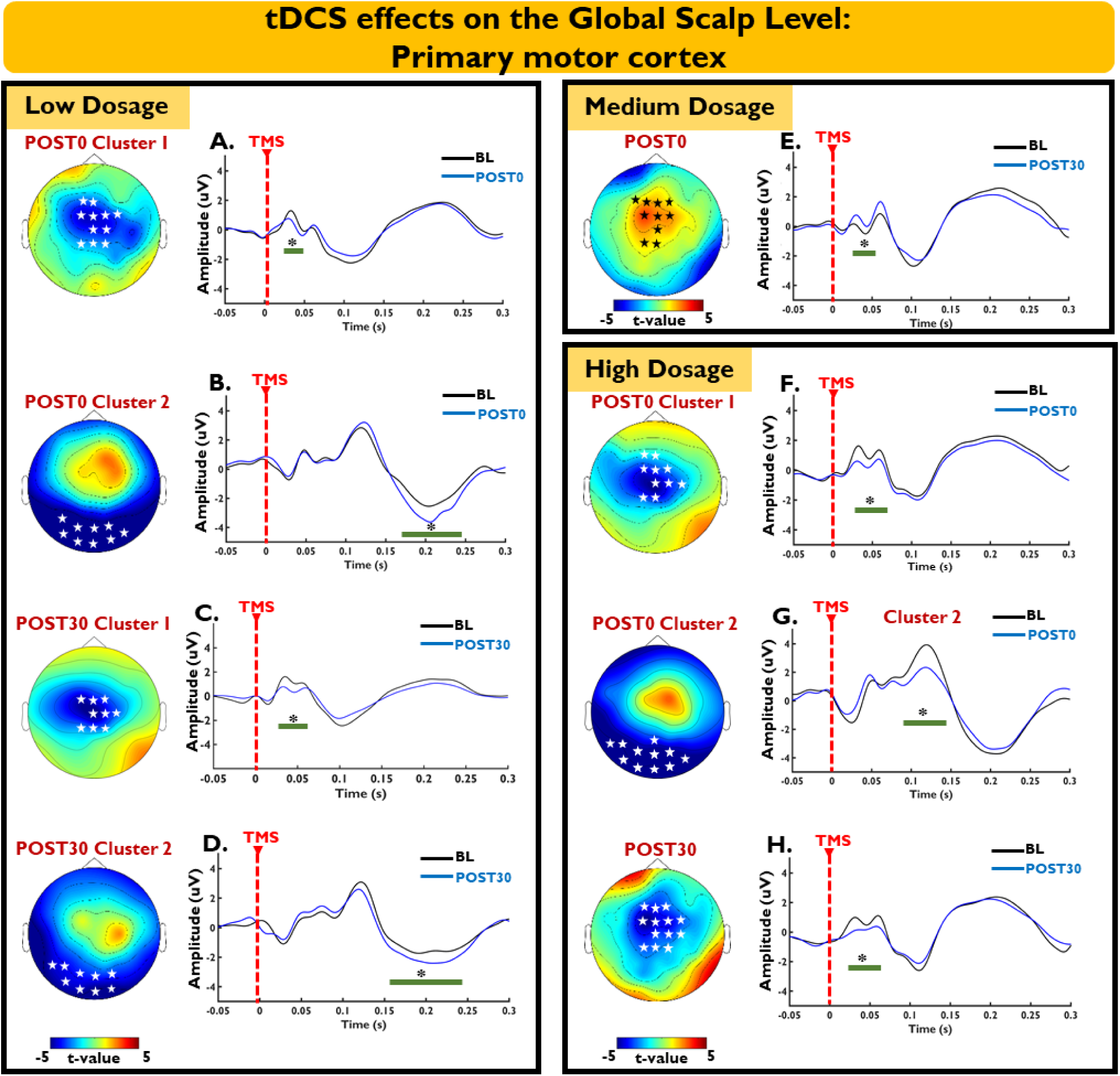
Global effects of tDCS on TMS-evoked Potentials (over the M1; comparison vs. baseline). The distributed effects of tDCS were evaluated via cluster-based permutation tests, immediately (POST0) to up to two hours after stimulation (POST30, POST60, POST120), over all electrodes. Topographic plots (distribution of the t-values) showing significant negative clusters (white stars) or positive clusters (black stars), together with the TEP deflections recorded over the EEG channels contribute to the significant clusters after low- (A-D), medium- (E) and high-dosage (F-H) tDCS over M1, and the green line represents the duration of the significant differences (*) between the respective time-points of tDCS-after effects vs. baseline. For further information regarding the specific electrodes forming each cluster refer to Figure 4-2. See also figure 4-1 for the global effects of tDCS on TMS-evoked Potentials (over M1; comparison vs. Sham).

For the distributed effects of ***tDCS over the PFC*,** the cluster-based permutation tests comparing the tDCS after-effects with respective baseline measures showed for medium-dosage tDCS a negative cluster over the fronto-central electrodes (POST0; p=0.024, time-period: 27-69 ms after TMS), a negative cluster over the fronto- central electrodes (POST30; p=0.036, time-period: 62-87 ms after TMS) and another negative cluster over the central electrodes (POST30; p=0.034, time-period: 79-111 ms after TMS). In addition, for high-dosage tDCS, a negative cluster was identified over the fronto-central electrodes (POST0; p=0.039, time-period: 27-72 ms after TMS) and over central electrodes (POST30; p=0.024, time-period: 21-60 ms after TMS), whereas a positive cluster was identified over a different group of central electrodes (POST30; p=0.026, time-period: 148-253 ms after TMS); Figure 5. Moreover, the results of the cluster-based permutation tests comparing the active tDCS conditions with sham measures showed a negative cluster for medium-dosage tDCS over the fronto-central electrodes (POST30; p=0.038, time-period: 23-69 ms after TMS); a negative cluster for high-dosage tDCS over the centro-medial electrodes (POST0; p=0.042, time-period: 34-86 ms after TMS) and another negative cluster over the right frontal electrodes for high-dosage tDCS (POST30; p=0.035, time-period: 25-96 ms after TMS); Figure 4-1.D-F.

**Figure 5.**
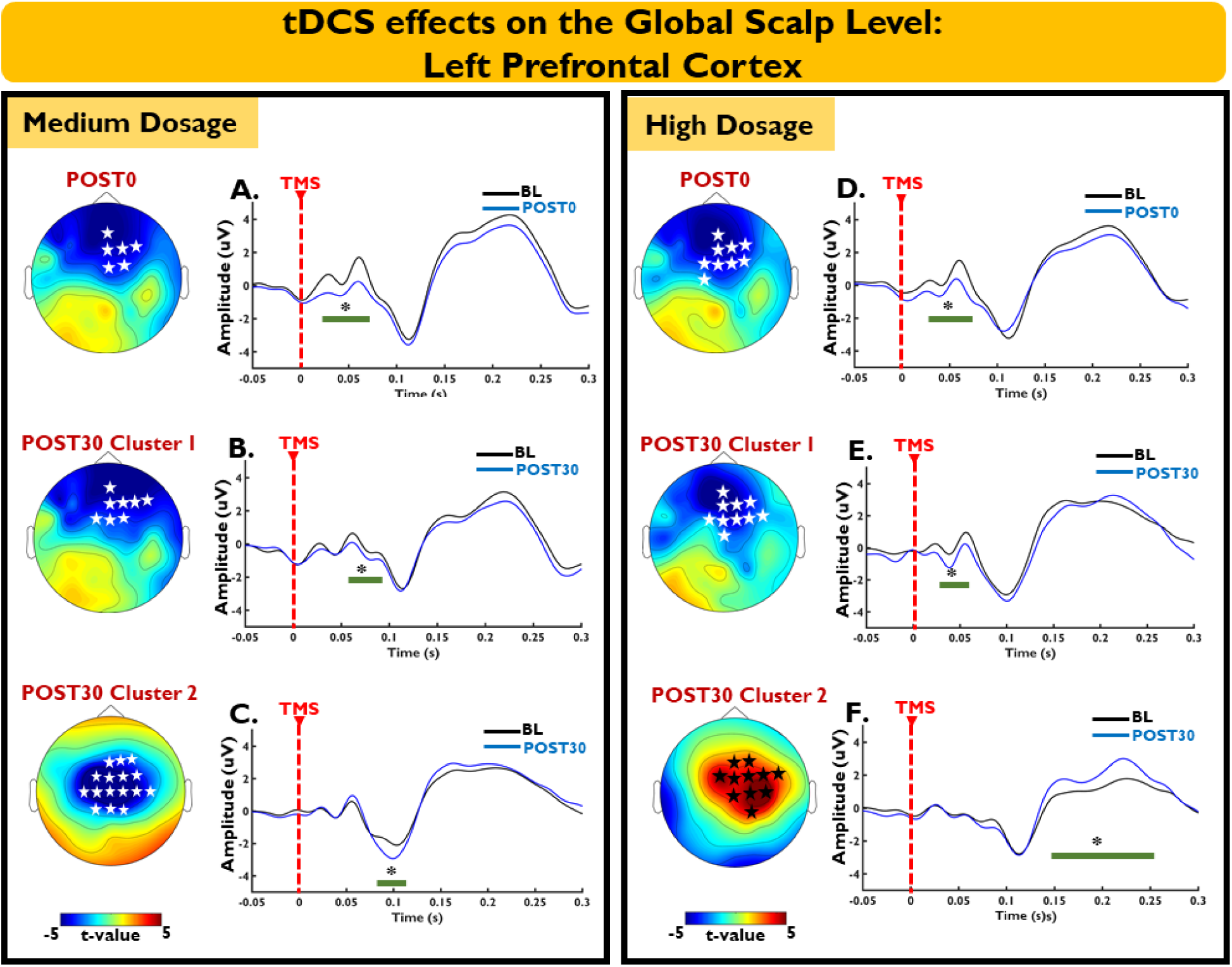
Global effects of tDCS on TMS-evoked Potentials (over the PFC; comparison vs. baseline). The distributed effects of tDCS were evaluated, using cluster based permutation test, immediately (POST0) to up to two hours after stimulation (POST30, POST60, POST120), over all electrodes. Topographic plots (distribution of the t-values) showing significant negative clusters (white stars) or positive clusters (black stars), together with the TEP deflections recorded over the EEG channels which constitute the significant cluster, after medium- (A-C), and high-dosage (D-F) tDCS over the prefrontal cortex. The green line represents the duration of the significant differences (*) between the respective time-points of tDCS-after effects vs. baseline. For further information regarding the specific electrodes forming each cluster refer to Figure 4-1. See also figure 4-1 for the global effects of tDCS on TMS-evoked Potentials (over PFC; comparison vs. Sham).

Together, these results indicate widespread effects of tDCS on TMS-evoked cortical reactivity at the global scalp level, which hints at the contribution of distant cortical networks on the overall tDCS efficacy.

#### 3.2.3. tDCS-altered TMS-evoked Oscillations (Regional Effects)

The primary rmANOVAs were conducted to test if the tDCS after-effects on TMS-evoked oscillations differed vs. baseline. For all tested frequency bands (θ, α, β, γ), rmANOVAs showed no significant main effects of condition, time-point, or their respective interactions (Table 3, Figure 6, Figure 7). However, significant results were observed between stimulation sites for all tested frequency bands (except Theta) and the interaction of ‘time-point × stimulation site’ (only for the Gamma frequency band), indicating a general lower power at the PFC in comparison with the M1 stimulation site. In addition, the secondary rmANOVAs, conducted to assess if the after-effects of the active tDCS protocols on TMS-evoked oscillations differed vs. sham values, showed no significant effect of neither the main factors condition, time-point, and stimulation sites, nor their interactions (Table 3-1, Figure 7-1).

**Figure 6.**
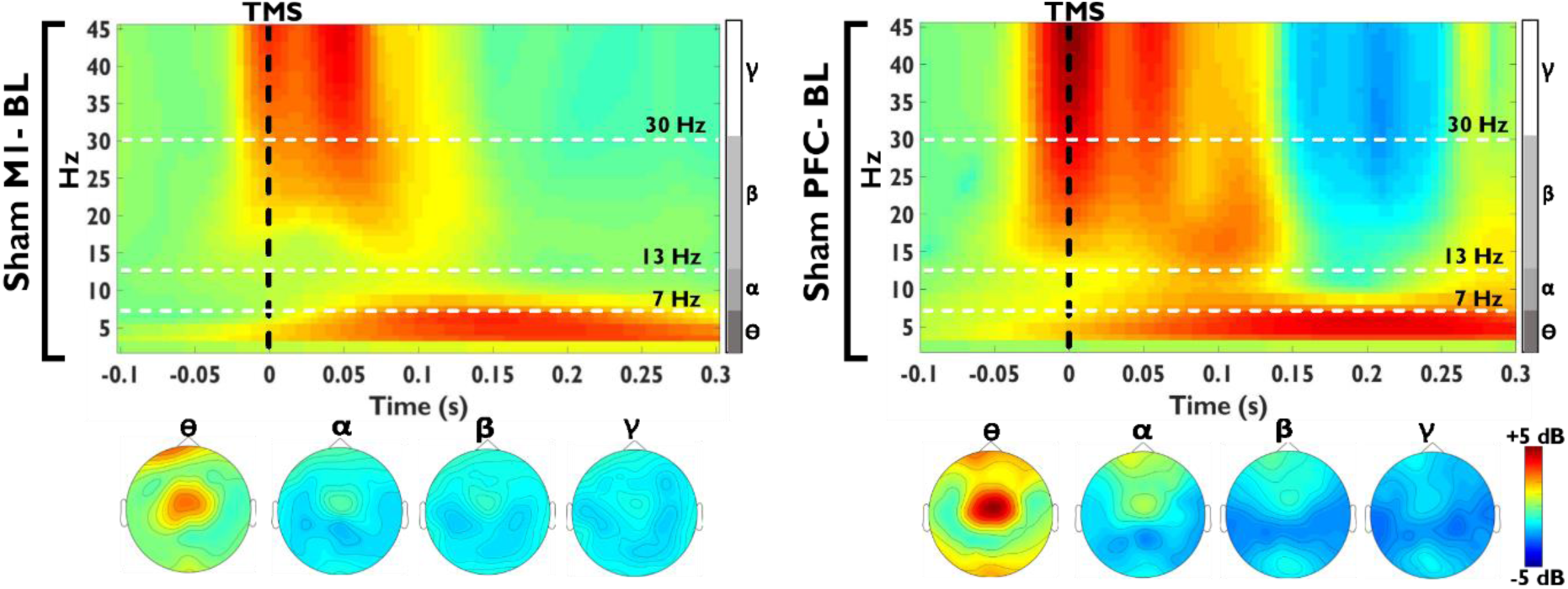
Local reperesentation of TMS-evoked Oscillations. Time-frequency representations (TFRs) of oscillatory power (Morlet wavelet; wavelet width: starting from 2.6 cycles and adding 0.2 cycle for each 1 Hz), normalized (db) to the respective baseline (−500 to −100ms), for ROI_M1_ (averaged FC1 and CP1 electrodes) and ROI_PFC_ (averaged FCz and Fz electrodes). The time-frequency plot shows the total power across different frequencies as a function of time following sham stimulation (baseline; BL) over the left primary motor cortex (ROI_M1_; **left**) and left DLPFC (ROI_PFC_; **right**). Power estimates were then calculated for four separate frequency bands, comprising Theta (θ; 4-7Hz), Alpha (α; 8-13Hz), Beta (β; 14-29Hz), and Gamma (γ; 30-45Hz), and a time window of 50–300ms. The topographic plots are showing the power estimates across four different frequency bands for the sham condition over M1 (left) or PFC (right) for the baseline measures; see Figure 6-1 for the TFR of all conditions and time-points. Note that the time-frequency and topographic plots are only illustrated for the baseline measures of sham conditions, as there were no significant differences of the regional tDCS after-effects on TMS-evoked oscillations in comparison with baseline and/or sham values, see figure 7, 7-1 for the calculated values across conditions and time-points.

**Figure 7.**
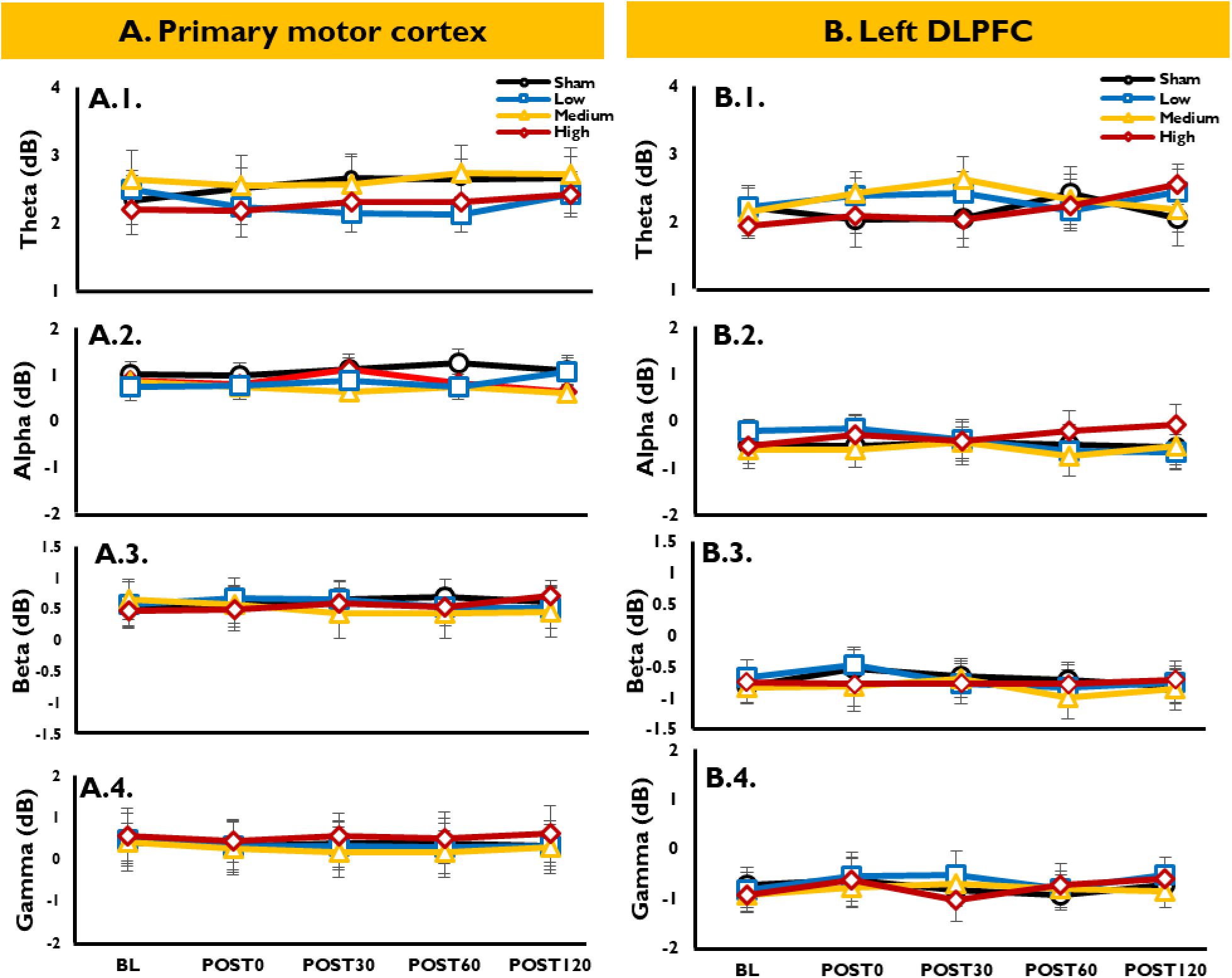
Local effects of tDCS on TMS-evoked oscillations (absolute values; comparison vs. baseline). Time-frequency representations (TFRs) of oscillatory power were calculated (Morlet wavelet; wavelet width: starting from 2.6 cycles and adding 0.2 cycle for each 1 Hz), and then normalized (dB) to the respective baseline (−500 to −100ms) for ROI_M1_ (averaged FC1 and CP1 electrodes) and ROI_PFC_ (averaged FCz and Fz electrodes). **A.1-4, B.1-4)** power estimates were then calculated before (BL) and at four time- points (immediately: POST0: POST30,POST60 and POST120) after tDCS, for four separate frequency bands, comprising Theta (θ; 4-7Hz), Alpha (α; 8-13Hz), Beta(β; 14-29Hz) and Gamma (γ; 30-45Hz), within a time window of 50–300ms. Error bars show the standard error of the mean (SEM). See figure 7-1 for the local effects of tDCS on TMS-evoked oscillations (normalized values).

**Table 3.**
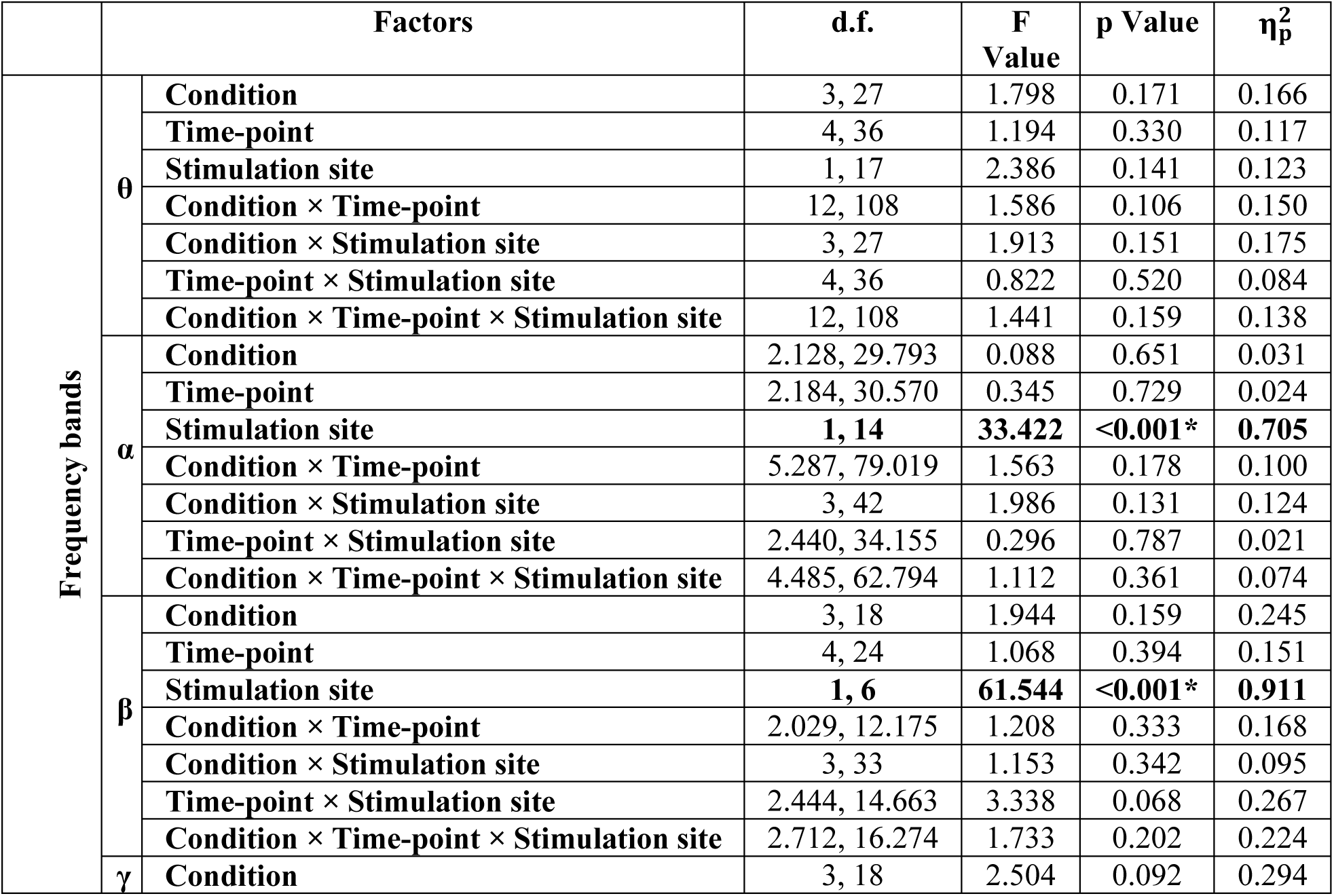
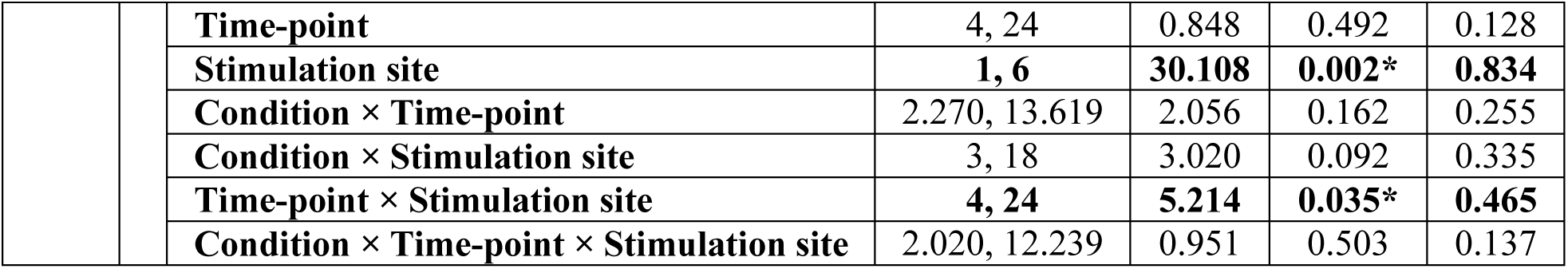
Results of the ANOVAs conducted for tDCS-induced alterations of cortical oscillations (absolute values). The primary rmANOVAs conducted to test the effects of tDCS on TMS-evoked oscillations vs. respective baseline values showed no significant effect of tDCS at the regional level on cortical oscillatory activities. However, significant results were observed between stimulation sites for all frequency bands (except Theta), and for the interaction ‘time- point × stimulation site’ (only for Gamma frequency bands), indicating a generally lower Gamma power of the PFC in comparison with the M1 stimulation site. d.f. = degrees of freedom, η*p*^2^ = partial eta squared. See also Table 3-1 for the results of the ANOVAs conducted for tDCS-induced alterations of cortical oscillations (normalized values).

| | Factors | | d.f. | F Value | p Value | $\eta_p^2$ |
| --- | --- | --- | --- | --- | --- | --- |
| Frequency bands | $\theta$ | Condition | 3, 27 | 1.798 | 0.171 | 0.166 |
|  |  | Time-point | 4, 36 | 1.194 | 0.330 | 0.117 |
|  |  | Stimulation site | 1, 17 | 2.386 | 0.141 | 0.123 |
| | | Condition × Time-point | 12, 108 | 1.586 | 0.106 | 0.150 |
| | | Condition × Stimulation site | 3, 27 | 1.913 | 0.151 | 0.175 |
| | | Time-point × Stimulation site | 4, 36 | 0.822 | 0.520 | 0.084 |
| | | Condition × Time-point × Stimulation site | 12, 108 | 1.441 | 0.159 | 0.138 |
| | $\alpha$ | Condition | 2.128, 29.793 | 0.088 | 0.651 | 0.031 |
|  |  | Time-point | 2.184, 30.570 | 0.345 | 0.729 | 0.024 |
|  |  | Stimulation site | **1, 14** | **33.422** | **<0.001\*** | **0.705** |
| | | Condition × Time-point | 5.287, 79.019 | 1.563 | 0.178 | 0.100 |
| | | Condition × Stimulation site | 3, 42 | 1.986 | 0.131 | 0.124 |
| | | Time-point × Stimulation site | 2.440, 34.155 | 0.296 | 0.787 | 0.021 |
| | | Condition × Time-point × Stimulation site | 4.485, 62.794 | 1.112 | 0.361 | 0.074 |
| | $\beta$ | Condition | 3, 18 | 1.944 | 0.159 | 0.245 |
|  |  | Time-point | 4, 24 | 1.068 | 0.394 | 0.151 |
|  |  | Stimulation site | **1, 6** | **61.544** | **<0.001\*** | **0.911** |
| | | Condition × Time-point | 2.029, 12.175 | 1.208 | 0.333 | 0.168 |
| | | Condition × Stimulation site | 3, 33 | 1.153 | 0.342 | 0.095 |
| | | Time-point × Stimulation site | 2.444, 14.663 | 3.338 | 0.068 | 0.267 |
| | | Condition × Time-point × Stimulation site | 2.712, 16.274 | 1.733 | 0.202 | 0.224 |
| | $\gamma$ | Condition | 3, 18 | 2.504 | 0.092 | 0.294 |

|  |  |  |  |  |  |
| --- | --- | --- | --- | --- | --- |
|  | **Time-point** | 4, 24 | 0.848 | 0.492 | 0.128 |
|  | **Stimulation site** | **1, 6** | **30.108** | **0.002\*** | **0.834** |
|  | **Condition × Time-point** | 2.270, 13.619 | 2.056 | 0.162 | 0.255 |
|  | **Condition × Stimulation site** | 3, 18 | 3.020 | 0.092 | 0.335 |
|  | **Time-point × Stimulation site** | **4, 24** | **5.214** | **0.035\*** | **0.465** |
|  | **Condition × Time-point × Stimulation site** | 2.020, 12.239 | 0.951 | 0.503 | 0.137 |

#### 3.2.4. tDCS-altered TMS-evoked Oscillations (global scalp Effects)

At the global scalp level, cluster-based permutation tests of TMS-evoked oscillations, comparing the tDCS after-effects with respective baseline measures showed for low-dosage tDCS over M1 a negative cluster over the occipito-parietal electrodes for the Alpha frequency band (α; POST30; p=0.021, time-period: 53-155 ms after TMS), a further negative cluster over the frontal electrodes for the Beta frequency band (β; POST30; p=0.019, time-period: 185-295 ms after TMS ), and two negative clusters for the Gamma frequency band over the fronto-central (γ; POST0, p= 0.012, time-period: 52-98 ms after TMS) and centro-parietal electrodes (γ; POST30, p=0.038, time-period 63-126 ms after TMS), Figure 8.A. For medium- dosage tDCS over M1, a positive cluster over the parietal electrodes (θ; POST30; p=0.024, time-period: 60-125 ms after TMS) and another positive cluster over the frontal electrodes (β; POST30; p=0.019, time-period: 185-295 ms after TMS) was observed, Figure 8.B. Furthermore, for high-dosage tDCS over M1, two negative clusters were identified over the occipital electrodes for the Theta frequency band, one at the POST0, and the other at the POST30 time-point (θ; POST0, p= 0.018, time-period: 65-118ms after TMS; POST30, p=0.034, time-period: 96-185 ms after TMS), as well as one negative cluster for the Alpha frequency band over the frontal electrodes (α; POST0, p= 0.018, time-period: 59-96 ms after TMS), two negative clusters for the Beta frequency band over the right fronto-central (β; POST0, p= 0.032, time-period: 54-83 ms after TMS), and right parietal electrodes (β; POST30, p=0.038, time-period 93-164 ms after TMS), and one negative cluster for the Gamma frequency band over the central electrodes (γ; POST30, p=0.022, time-period 87-196 ms after TMS), Figure 8.C. No significant clusters were however identified between sham measures and active M1 tDCS dosages.

**Figure 8.**
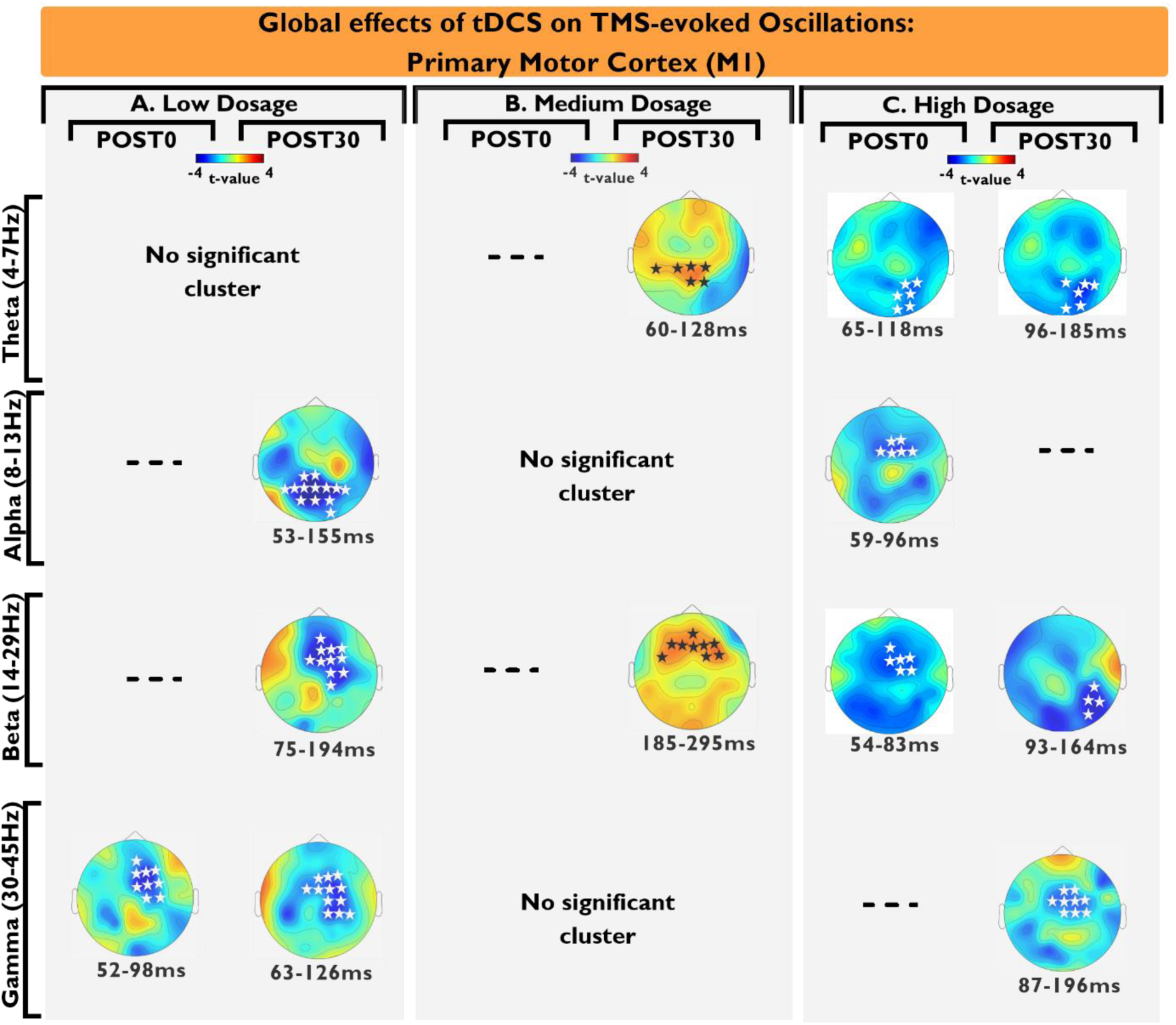
Global effects of tDCS on TMS-evoked oscillations (tDCS over M1; comparison vs. baseline). The distributed effects of tDCS on oscillatory brain activity were evaluated by cluster-based permutation tests, immediately (POST0) to up to two hours after stimulation (POST30, POST60, POST120), over all electrodes. Topographic plots (distribution of the t-values) show significant negative clusters (white stars) or positive clusters (black stars), over the EEG channels, which constitute significant clusters after low- (A), medium- (B), and high-dosage (C) tDCS for Theta, Alpha, Beta and Gamma frequency bands. The permutation tests were calculated within the time window of 50-300ms after TMS. Significant clusters were identified only for POST0 and/or POST30. Dashed lines indicate no significant cluster at the respective time point. The duration below each topographic plot indicates the time period of the significant clusters. Further detailed information regarding the specific electrodes forming each cluster see Figure 8-2.

For the distributed effects of ***tDCS over the PFC*,** the cluster-based permutation tests comparing the tDCS after-effects with respective baseline measures showed for medium-dosage tDCS a negative cluster over the left centro-parietal electrodes for the Theta frequency band (θ; POST30; p=0.031, time-period: 51-108 ms after TMS), a further negative cluster over the frontal electrodes for the Alpha frequency band (α; POST30; p=0.018, time-period: 75-213 ms after TMS), two negative clusters for the Beta frequency band over the right centro-frontal electrodes (β; POST0, p=0.042, time-period 62-93 ms after TMS and POST30, p=0.022, time-period 71-131 ms after TMS), as well as one negative cluster over the right frontal electrodes for the Gamma frequency band (γ; POST30; p=0.014, time-period: 99-288 ms after TMS), Figure 9.B. For high-dosage tDCS, a negative cluster was identified over the right centro-frontal electrodes for the Alpha frequency band (α; POST0; p=0.023, time-period: 89-201 ms after TMS), and two negative clusters for the Gamma frequency band over the fronto-central electrodes (γ; POST0; p=0.029, time-period: 87-196 ms after TMS; POST30; p=0.003, time-period: 67-116 ms after TMS), Figure 9.C. No significant clusters were however identified between sham measures and active PFC tDCS dosages.

**Figure 9.**
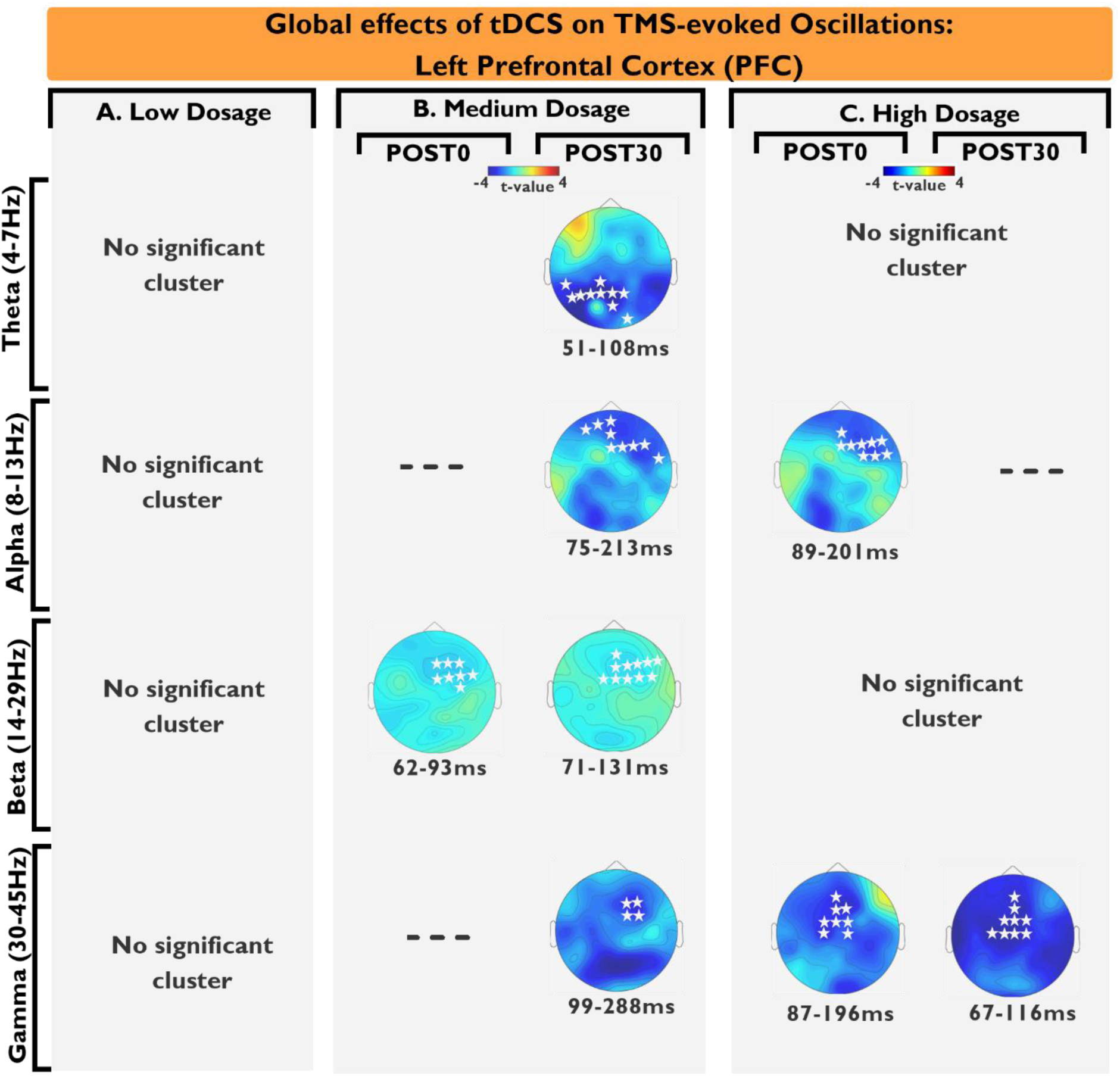
Global effects of tDCS on TMS-evoked Oscillations (over the PFC; comparison vs. baseline). The distributed effects of tDCS on oscillatory brain activity were evaluated by cluster-based permutation tests, immediately (POST0) to up to two hours after stimulation (POST30, POST60, POST120), over all electrodes. Topographic plots showing significant negative clusters (white stars) or positive clusters (black stars), over the EEG channels which constitute the significant clusters after low- (A), medium- (B), and high-dosage (C) tDCS for Theta, Alpha, Beta, and Gamma frequency bands. The permutation tests were calculated for a time window of 50-300ms after TMS. Significant clusters were identified only after POST0 and/or POST30. Dashed lines indicate no significant cluster at the respective time point. The duration below each topographic plot indicates the time period of the significant clusters. For further detailed information regarding the specific electrodes forming each cluster refer to Figure 8-2.

Together, these results indicate widespread effects of tDCS on TMS-evoked oscillations at the global scalp level, which hints at the contribution of distant cortical networks to the overall tDCS effects.

#### 3.2.5. tDCS-altered TMS-elicited MEPs

The 2-factorial ANOVA (condition-4 levels, and time point-5 levels), conducted for the MEP amplitudes (absolute values) revealed significant main effects of time- point, and a significant ‘condition × time-point’ interaction (*F*_(1.550,26.534)_ = 70.463, *p* < 0.001, ɳ^2^ = 0.806). Post hoc tests comparing tDCS after-effects with respective baseline values revealed a significant MEP amplitude reduction after low- (POST0, POST30) and high-dosage (POST0, POST30 and POST60) tDCS (Table 4.A Figure 10). In addition, the secondary rmANOVAs on ΔMEPs (normalized values and excluding baseline measures), revealed significant effects of the main factors condition and time-point and their respective interactions (*F*_(2.152,36.586)_ = 3.094, *p* = 0.002, ɳ^2^ = 0.154) (Table 4.B, Figure 10-1). Post hoc tests comparing after-effects of real with sham stimulation showed ΔMEP amplitude reductions after low- and high-dosage (both at POST0, POST30) tDCS (Figure 10-1, Table 4.B). In summary, the results show dosage-dependent effects of tDCS on MEP amplitude alterations, with low- and high-dosage tDCS resulting in a cortico-spinal excitability reduction, while medium-dosage tDCS had no effects.

**Table 4.** Results of the ANOVAs conducted for evaluation of tDCS-generated alterations of TMS-evoked MEPs. The primary rmANOVA (**A**; absolute values) results show a significant main effect of time-point, and a significant ‘condition × time-point’ interaction. In addition, the secondary rmANOVA MEPs (**B**; normalized values, baseline measures excluded from the analysis) showed significant effects of the main factors condition and time-point, and their respective interactions. Asterisks indicate significant effects (p < .05), d.f. = degrees of freedom, η*p*^2^ = partial eta squared.

| | Factors | | d.f. | F Value | p Value | $\eta_p^2$ |
| --- | --- | --- | --- | --- | --- | --- |
| A | MEP | Condition | 3, 51 | 1.212 | 0.315 | 0.067 |
|  |  | Time-point | 1.599, 27.157 | 78.668 | <0.001* | 0.822 |
|  |  | Condition × Time-point | 1.550, 26.534 | 70.463 | <0.001* | 0.806 |
| B | ΔMEP | Condition | 3, 51 | 13.547 | <0.001* | 0.443 |
|  |  | Time-point | 1.586, 26.968 | 3.155 | 0.033* | 0.157 |
|  |  | Condition × Time-point | 2.152, 36.586 | 3.094 | 0.002* | 0.154 |

**Figure 10.**
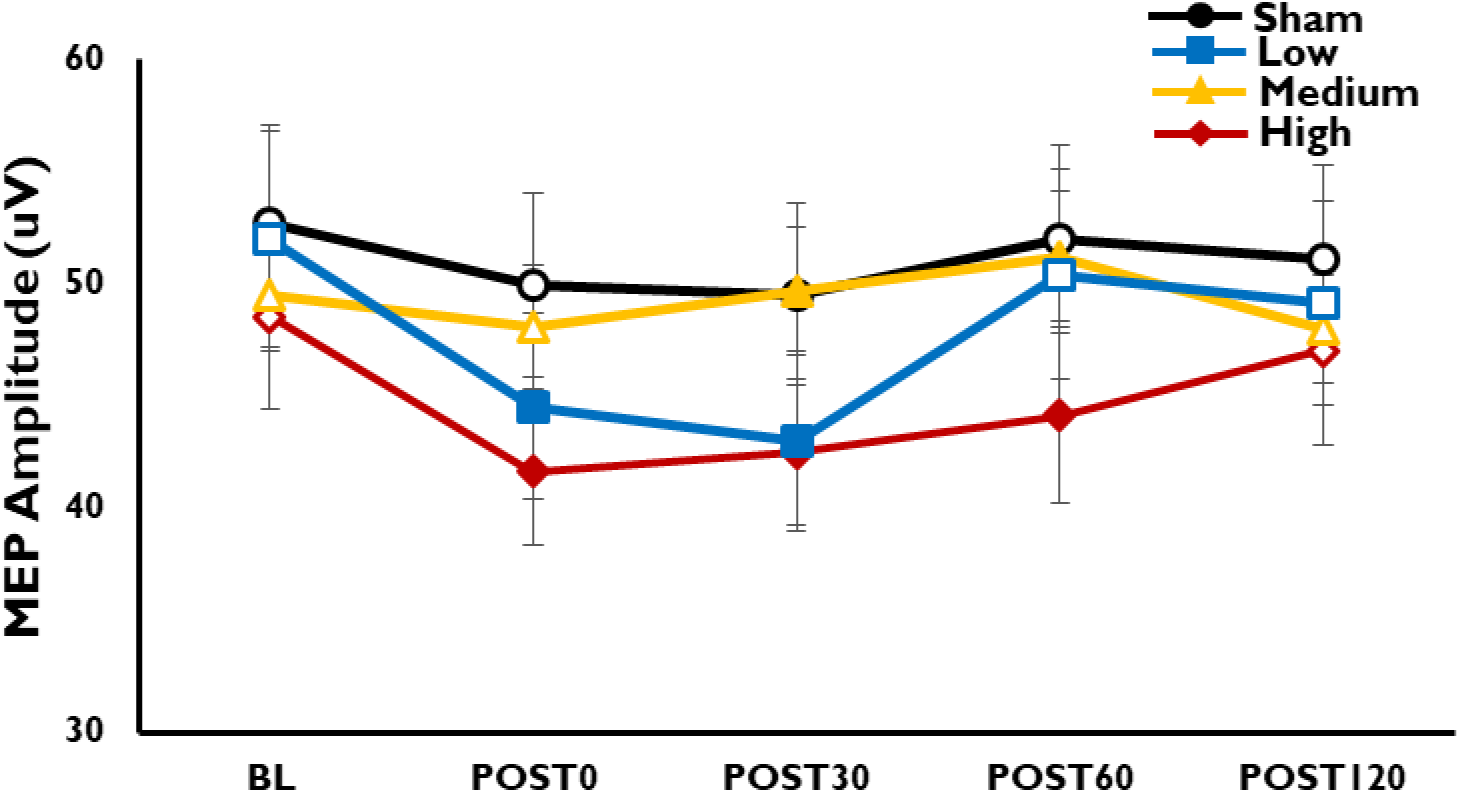
Local effects of tDCS on TMS-elicited MEPs (absolute values). Low, medium-, and high- cathodal tDCS intensities, and sham tDCS, were applied over the primary motor cortex (M1). The tDCS effects on cortico-spinal excitability were then evaluated, immediately (POST0) to up to two hours after stimulation (POST30, POST60, POST120). Filled symbols show significant differences vs. baseline. Error bars show the standard error of the mean (SEM). For the comparison of the tDCS effects vs. sham and/or between active conditions see figure 10-1.

### 3.3. Association between tDCS-altered TMS-evoked Potentials and TMS-elicited MEPs and tDCS-induced electrical fields

The Pearson correlation results indicate a significant positive relation between P30 and MEP for low- (POST0: r = 0.56, p = 0.031), and high-dosage tDCS (POST0: r = 0.53, p = 0.037) (Figure 11.A). No significant correlations were found between other TEPs and MEPs across conditions and time-points (all with p > 0.05). There was a significant positive correlation between tDCS-induced EFs and MEP amplitude changes after high-dosage tDCS (POST30: r = 0.51, p = 0.029), indicating that subjects with higher EFs over the targeted area showed larger MEP amplitude changes (Figure 11.B). However, the correlation results did not show any significant relationship between tDCS-induced EFs and tDCS-altered TEP peaks (all with p > 0.05). Moreover, the tDCS-induced EFs over the PFC were significantly lower than the tDCS-induced EF over M1 (p=0.011), Figure 11.C. Interestingly, tDCS-induced EFs over M1 and the PFC did not correlate significantly (r=0.31, p=0.074). Note that the tDCS-induced EFs were calculated using a GM mask over the TMS-induced effective EFs at each stimulation site (see respective section of the method).

**Figure 11.**
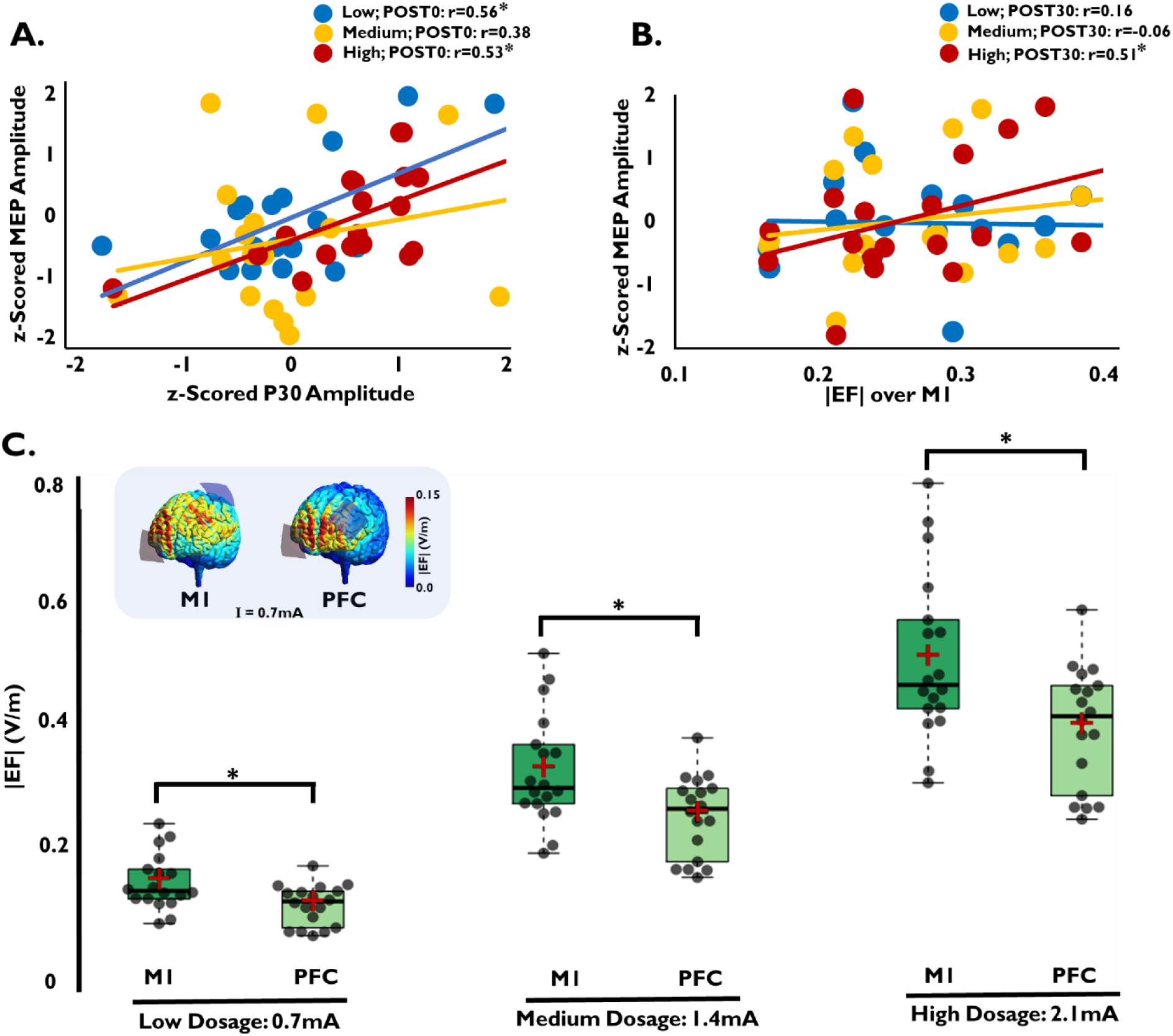
Association between tDCS-generated TMS-evoked Potential and TMS-elicited MEP alterations and tDCS-induced electrical fields. Pearson correlations were calculated to test the relationship between tDCS-generated TEP and MEP alterations (A), tDCS-generated MEP alterations and tDCS-induced EF (B). In addition, the differences between the tDCS-induced EFs (strength |EF|) over the M1 and PFC were explored via paired Student’s t-test (C: Box plots (whiskers extend to minimum and maximum individual values (shown in dark gray circles)); also the median (horizontal black line in the box), and means (shown as red + symbols) of the EFs are shown). Note that the estimated tDCS-induced EFs were calculated for low- (0.7mA), medium- (1.4mA), and high- intensity (2.1mA) tDCS, but that the results can linearly be scaled for the other tDCS intensities based on a single intensity according to the quasi-static approximation (Miranda et al., 2013). The colors of the anatomical pictures are illustrating EF magnitudes induced by tDCS (I=0.7mA) estimated via SimNIBS open-source software with its default parameters and head model (ernie.msh); for individual EF maps see Figure 11-1 and Figure 11-2. Asterisks show significant differences.

In summary, the results showed an association between the early TEP peaks (only P30) and MEPs, which is in line with previous findings (Mäki and Ilmoniemi, 2010). However, this was not consistent across all active conditions and for all respective time-points of tDCS-after-effects. Also, the regional tDCS-induced EF over the PFC is, on average and across participants, lower than that over M1, but this might not be identical at the individual level in each case.

### 3.4. Qualitative assessment of tDCS protocols

The results of the Chi-square tests indicate no significant heterogeneity for any of the tDCS dosages (sham: χ2=0.111, p=0.739; low-dosage: χ2=2.778, p=0.096; medium-dosage: χ2=1.778, p=0.182; high-dosage: χ2=1.000, p=0.317), which shows successful blinding. Also, the ANOVAs conducted for the side-effects showed no significant effects either during or 24h after stimulation (Table 5). Guesses of received stimulation intensity vs actual intensity are shown in Table 5-1. Ratings of the presence and intensity of side-effects are documented in Table 5-2.

**Table 5:** The presence and intensity of side effects were analyzed by one-way repeated-measure ANOVAs. No significant effects of side effects were identified either during or 24h after stimulation. d.f.= degrees of freedom. η*p*^2^ = partial eta squared. Guesses of received stimulation intensity vs actual intensity are shown in Table 5-1. Ratings of the presence and intensity of side-effects are documented in Table 5-2.

| | Side-effects | d.f | F Value | $\eta_p^2$ | p Value |
| --- | --- | --- | --- | --- | --- |
| **During stimulation** | **Visual Phenomena** | 7, 119 | 1.567 | 0.084 | 0.152 |
|  | **Itching** | 7, 119 | 0.765 | 0.043 | 0.618 |
|  | **Tingling** | 7, 119 | 1.780 | 0.095 | 0.098 |
|  | **Pain** | 3.106, 52.800 | 1.041 | 0.058 | 0.384 |
| **24 hours after stimulation** | **Redness** | 7, 119 | 0.363 | 0.021 | 0.922 |
|  | **Headache** | 2.432, 41.342 | 1.157 | 0.064 | 0.332 |
|  | **Fatigue** | 7, 119 | 1.560 | 0.084 | 0.154 |
|  | **Concentration** | 3.038, 51.650 | 1.129 | 0.062 | 0.346 |
|  | **Nervousness** | 3.100, 52.706 | 2.209 | 0.115 | 0.096 |
|  | **Sleep Problem** | 7, 119 | 1.630 | 0.087 | 0.133 |

## 4. Discussion

In this study, we explored the dosage-dependent neurophysiological effects of cathodal tDCS at two targeted stimulation sites, the M1 and PFC. In a single-blind and sham-controlled repeated measures design, four cathodal tDCS dosages of low, medium, high intensity, and sham stimulation were applied over the M1 and PFC. The after-effects were then tested by TMS-EEG and TMS-MEP techniques, at the regional and global scalp level, for TMS-evoked cortical reactivity, oscillations, and MEP amplitude alterations. In general, at the regional level, we observed a nonlinear dosage-dependent effect of M1 tDCS for TMS-evoked early positive TEPs, and MEPs, whereas PFC tDCS decreased almost uniformly the early positive TEP peaks; we however did not observe regional after-effects of tDCS on TMS-evoked oscillations. In addition, at the global scalp level, we observed distributed effects of tDCS on both, TMS-evoked potentials and oscillations. Computational modeling of tDCS-induced EFs showed relatively lower EFs over the PFC in comparison with M1 with the respective electrode positions, which was reflected by relatively smaller physiological effects after PFC, as compared to M1 tDCS. Blinding was successful, and all participants tolerated tDCS well. These findings are discussed in detail below.

### 4.1. Effects of tDCS on TMS-evoked Potentials, MEPs and Oscillations (regional effects)

Previous TMS-EEG studies have suggested that the early P30 TEP peak might reflect excitatory processes at the stimulation site (Ilmoniemi et al., 1997; Rogasch and Fitzgerald, 2013; Darmani and Ziemann, 2019; Rogasch et al., 2020). In addition, recent studies of our group have shown dosage-dependent non-linear cathodal ***tDCS after-effects over M1***, which are suggested to be linked with calcium channel dynamics (Nitsche et al., 2004; Mosayebi-Samani et al., 2020). These together might explain, at the regional level, the observed reduction of the P30 TEP peak after low (which is in line with previous studies (Pellicciari et al., 2013; Ahn and Fröhlich, 2021)) and high dosage tDCS over M1, and an enhancement after medium dosage tDCS, which is also in accordance with the tDCS after-effects on MEP amplitude changes. Likewise, the alteration of the P60 peak, which is suggested to be controlled by fast glutamatergic neurotransmission in the stimulated cortical network (Ferreri et al., 2011; Rogasch and Fitzgerald, 2013; Cash et al., 2017; Darmani and Ziemann, 2019; König et al., 2019), might be caused by the dependency of cathodal tDCS after-effects from glutamate, specifically the reduction of glutamate after low intensity cathodal tDCS, and the presumably glutamate-dependent calcium dynamics causing the after-effects of medium- and high-dosage tDCS (Nitsche et al., 2003; Nitsche et al., 2004; Stagg et al., 2009; Mosayebi-Samani et al., 2020), as the effects are equivalent to the P30 peak. However, alternative explanations cannot be ruled out, e.g. antagonistic responses of different cortical layers to an identical stimulation dosage (Purpura and McMurtry, 1965; Sun et al., 2020). At present, these explanations are speculative and should be explored directly in future studies.

A null effect of tDCS on late-latency peaks (at periods ∼100 ms, and ∼200 ms; e.g. N100 and P200) was observed. Previous studies have linked these peaks to changes of local GABA-related activity (Connors et al., 1988; Ferreri et al., 2011; Premoli et al., 2014b), as well as interhemispheric excitatory-inhibitory activity (Nikulin et al., 2003; Bender et al., 2005; Voineskos et al., 2010; Rogasch and Fitzgerald, 2013; Premoli et al., 2014b; Du et al., 2017; Darmani and Ziemann, 2019). Therefore, it might be speculated that local GABA reduction, which would decrease this potential, would be counteracted upon by glutamate reduction-related enhancement of the N100, which would then result in a zero-net effect of tDCS on the N100. Sensory and somatosensory evoked potentials might also contribute to these late-latency peaks resulting from the TMS-elicited clicking noise and coil vibration (Gordon et al., 2018a; Biabani et al., 2019; Conde et al., 2019; Ahn and Fröhlich, 2021; Biabani et al., 2021), and diminish effect differences between sham and real stimulation conditions. Knowledge about the neurophysiological foundations of TEPs is however still rudimentary and warrants further investigation.

The same mechanisms as outlined above might also contribute to the after-effects of ***tDCS over the PFC***, but anatomical and/or neurophysiological differences might contribute to the gradual differences of the effects with respect to both cortical areas. The lower inter-electrode distance and larger scalp-to-cortex distance in case of prefrontal stimulation (Kozel et al., 2000; McConnell et al., 2001) resulted in lower tDCS-induced EFs over the PFC, as shown by computational results (Faria et al., 2011; Laakso et al., 2015; Opitz et al., 2015; Mosayebi-Samani et al., 2021), and, therefore, potentially explains the observed uniform reduction of P30 and P60 after different dosages of tDCS over the PFC, in comparison with the observed dosage- dependent non-linear effects of tDCS over M1. Moreover, pharmacological studies showed that dopamine, which is more prevalent in the prefrontal, as compared to the motor cortex (Gaspar et al., 1995), strengthens the reduction of cathodal tDCS- generated cortical excitability diminution (Kuo et al., 2008), and might thus have prevented the conversion effects of medium-dose tDCS into cortical excitability enhancement, as observed for the P30 amplitude after tDCS over M1.

The results, however, did not show any significant effects of tDCS on the regional TMS-evoked oscillations over both stimulation sites, which is in line with findings of other NIBS protocols (Chung et al., 2017).

### 4.2. Effects of tDCS on TMS-evoked Potentials and Oscillations (Global scalp effects)

At the global scalp level, widespread effects across multiple EEG channels were observed for both, tDCS effects on TMS-evoked potentials and oscillations. This is in line with our computational modelling results showing spreading of tDCS- induced EFs from the stimulation site across remote regions, but also with previous findings, showing modulation of activity across functional cortical networks, as measured by EEG (Polanía et al., 2011; Cosmo et al., 2015), and other neuroimaging techniques (Lang et al., 2005; Keeser et al., 2011). Interestingly, while we did not observe regional effects of tDCS for the late TEPs, the clusters revealed in the global scalp analysis suggest a contribution of these late-latency peaks to the distributed network which might be related to tDCS-altered GABA activity (Premoli et al., 2014b; Premoli et al., 2014a), additional to confounding factors explained above. These late-latency changes might also be secondary responses that reflect the regional effects at the stimulation site due to effective connectivity.

In addition, tDCS after-effects were observed on TMS-evoked oscillations across different frequency bands. Previous studies have suggested the β–band to be associated with motor (Zhang et al., 2008), but also cognitive processes (Buschman and Miller, 2007), and γ band oscillations are known to play a role in neuronal communication across cortical regions, as well as integration of sensory information (Howard et al., 2003; Womelsdorf et al., 2007). With this in mind, GABA has been shown to play a role in modulation of γ and β activity (Bartos et al., 2007; Gaetz et al., 2011), and the observed changes in the present study might likely be explained by the dependency of the tDCS after-effect on GABA modulation (Stagg et al. 2009). Given the mixed results of tDCS effects on neural oscillations, as seen in this study and the relevant literature (Pellicciari et al., 2013; Hill et al., 2017; Gordon et al., 2018b; Ahn and Fröhlich, 2021), more research is clearly needed to better define any specific tDCS effects on TMS-evoked neural oscillations, and respective underlying mechanisms. Beyond these physiological considerations, however, it should be recognized that volume conduction effects cannot be ruled out (van den Broek et al., 1998).

### 4.3. Limitations and future directions

This study should be interpreted within the context of some limitations. First, this was an exploratory study. Data were acquired from a relatively small sample over a couple of sessions involving different interventions. Thus, the results are preliminary and should be replicated in follow-up studies with larger samples. In addition, due to a technical issue related to the used navigation system in the current study, an EEG 10-10 based method was selected for the identification of the left DLPFC target (F3) (Herwig et al., 2003; Hill et al., 2017). TMS coil positioning based on neuronavigation with MNI coordinates would have increased exactness and should be preferred in future studies (Rusjan et al., 2010; Fitzgerald, 2021). Furthermore, late-latency peaks (about 100 ms after the TMS pulse) might reflect also non-direct effects of TMS due to the contribution of auditory-evoked potentials resulting from the TMS clicking noise and bone-conducted sensory responses caused by coil vibration (Gordon et al., 2018a; Biabani et al., 2019; Conde et al., 2019; Ahn and Fröhlich, 2021; Biabani et al., 2021). A recent study has suggested methods to reduce these confounding factors (Rocchi et al., 2021), which should be considered in future studies. In the present study, we did not use a thin layer of foam under the TMS coil, which could decrease the non-direct effects of TMS because of coil vibration, because this has the disadvantage of increasing RMT, and thus TMS intensity required for excitability measures because of the resulting larger coil to brain distance. This approach should however be considered in future studies. We have adjusted the TMS intensity to 100% of RMT as a compromise to receive reasonable TEP, but also reduce indirect TMS effects. For obtaining MEP, this stimulation intensity is however comparatively low and might have affected respective measures.

Moreover, in the current study, we used relatively large tDCS electrodes to evaluate the effect of conventional tDCS protocols. Several EEG electrodes over the targeted areas had, therefore, to be removed, which limits the selection of relevant EEG electrodes for the analysis of regional effects. Future studies might consider smaller and/or other relevant electrode montages, introduced in recent studies (Romero Lauro et al., 2014; Hill et al., 2017; Gordon et al., 2018b). The selection of tDCS protocols in the current study was based on our foregoing study results of tDCS over M1 (Mosayebi Samani et al., 2019). We chose the protocols based on those which were most promising to reveal dosage-dependent non-linear effects, and thus our choice was effect-based. It included the lowest dosage used to induce an excitability diminution applied in that study (lowest intensity, and duration (15min)), a medium dosage, which induced excitability-enhancing effects (tDCS for 20min, medium intensity), and a high-dosage, which induced excitability-diminishing effects (tDCS for 20min; high intensity). While blinding was efficient with respect to stimulation intensity, we cannot be sure that blinding was affected by stimulation duration, however identical stimulation durations were associated with different intensities.

Also, previous studies showed a relatively uniform enhancement of motor cortical excitability following different anodal tDCS dosages (Jamil et al., 2017; Agboada et al., 2019). Therefore, the results of the current study might not directly translate to anodal stimulation effects, which needs further investigation. Likewise, pharmacological and neuroimaging studies highlighted polarity-dependent effects of tDCS on GABA and/or glutamatergic activities, among other receptors/neurotransmitters (Nitsche et al., 2003; Stagg et al., 2009), which might link to the TEP peak alterations (as outlined above). The impact of tDCS on positive or negative TEPs might furthermore depend on the orientation of the tDCS-induced electrical field. To explore this, however, also anodal tDCS would have been required, which was out of the scope of this study, and need further investigations. Moreover, the results of this study might not be one-to-one transferable to other cortical areas, other populations (Ghasemian-Shirvan et al., 2020; Ghasemian-Shirvan et al., 2022), as well as behavioral tDCS studies. Finally, the different effects obtained by prefrontal and motor cortex stimulation, as identified in this study, should be carefully evaluated in future studies, as it is not clear if these are due to biological differences between respective areas, or different current densities at the cortical level, due to anatomical differences. Further work, including computational modeling, might help to clarify this issue.

## Conclusion

The results of this study show neurophysiological effects of motor cortex tDCS at the regional and global scalp level on TMS-evoked cortical reactivity, which are comparable with respective cortico-spinal excitability effects, measured by TMS-generated MEP. Low- and high-dosage motor cortex tDCS reduced the early positive TEP peak and MEP amplitudes, whereas an amplitude enhancement was observed for the medium dosage of motor cortex tDCS. In contrast, prefrontal low-, medium- and high-dosage tDCS almost uniformly reduced the early positive TEP peak amplitudes. Furthermore, over both cortical areas, regional modulatory effects of tDCS were not observed for late TEP, and TMS-evoked oscillations. However, at the global scalp level, the results suggest a distributed effect of tDCS on both, TMS-evoked potentials and oscillations. The specific differences of the effects of tDCS might be related to physiological, anatomical, and receptor and transmitter differences of motor and non-motor areas. The overall results provide the first direct comparison of tDCS effects over different brain areas at the physiological level, which will further consolidate the rationale for extending tDCS applications at both basic and clinical levels.

## Supporting information

Supplemental TablesFigures

## Acknowledgment

We thank Prof. Marcello Massimini, Dr. Silvia Casarotto, and Dr. Matteo Fecchio for their important comments on the manuscript, and Dr. Asif Jamil, Dr. Fatemeh Yavari, Tobias Blanke, and Nina Abich for their valuable support during the study.

## Reference

Agboada D, Mosayebi Samani M, Jamil A, Kuo M-F, Nitsche MA (2019) Expanding the parameter space of anodal transcranial direct current stimulation of the primary motor cortex. Scientific reports 9:18185.

Ahn S, Fröhlich F (2021) Pinging the brain with transcranial magnetic stimulation reveals cortical reactivity in time and space. Brain stimulation 14:304–315.

Antal A, Kincses TZ, Nitsche MA, Bartfai O, Paulus W (2004) Excitability changes induced in the human primary visual cortex by transcranial direct current stimulation: direct electrophysiological evidence. Investigative ophthalmology & visual science 45:702–707.

Awiszus F, Borckardt J (2012) TMS Motor Threshold Assessment Tool 2.0 2012. Available from: http://clinicalresearcher. org/software. htm [accessed 19.10. 2012].

Bartos M, Vida I, Jonas P (2007) Synaptic mechanisms of synchronized gamma oscillations in inhibitory interneuron networks. Nature Reviews Neuroscience 8:45–56.

Beam W, Borckardt JJ, Reeves ST, George MS (2009) An efficient and accurate new method for locating the F3 position for prefrontal TMS applications. Brain stimulation 2:50–54.

Bender S, Basseler K, Sebastian I, Resch F, Kammer T, Oelkers-Ax R, Weisbrod M (2005) Electroencephalographic response to transcranial magnetic stimulation in children: evidence for giant inhibitory potentials. Annals of neurology 58:58–67.

Biabani M, Fornito A, Mutanen TP, Morrow J, Rogasch NC (2019) Characterizing and minimizing the contribution of sensory inputs to TMS-evoked potentials. Brain stimulation 12:1537–1552.

Biabani M, Fornito A, Coxon JP, Fulcher BD, Rogasch NC (2021) The correspondence between EMG and EEG measures of changes in cortical excitability following transcranial magnetic stimulation. The Journal of physiology 599:2907–2932.

Bikson M, Grossman P, Thomas C, Zannou AL, Jiang J, Adnan T, Mourdoukoutas AP, Kronberg G, Truong D, Boggio P (2016) Safety of transcranial direct current stimulation: evidence based update 2016. Brain stimulation 9:641–661.

Brunoni AR, Amadera J, Berbel B, Volz MS, Rizzerio BG, Fregni F (2011) A systematic review on reporting and assessment of adverse effects associated with transcranial direct current stimulation. Int J Neuropsychopharmacol 14:1133–1145.

Buschman TJ, Miller EK (2007) Top-down versus bottom-up control of attention in the prefrontal and posterior parietal cortices. Science 315:1860–1862.

Carbunaru R, Durand DM (2001) Toroidal coil models for transcutaneous magnetic simulation of nerves. IEEE Transactions on Biomedical Engineering 48:434–441.

Cash RFH, Noda Y, Zomorrodi R, Radhu N, Farzan F, Rajji TK, Fitzgerald PB, Chen R, Daskalakis ZJ, Blumberger DM (2017) Characterization of Glutamatergic and GABAA-Mediated Neurotransmission in Motor and Dorsolateral Prefrontal Cortex Using Paired-Pulse TMS–EEG. Neuropsychopharmacology : official publication of the American College of Neuropsychopharmacology 42:502–511.

Chung SW, Lewis BP, Rogasch NC, Saeki T, Thomson RH, Hoy KE, Bailey NW, Fitzgerald PB (2017) Demonstration of short-term plasticity in the dorsolateral prefrontal cortex with theta burst stimulation: A TMS-EEG study. Clinical neurophysiology : official journal of the International Federation of Clinical Neurophysiology 128:1117–1126.

Conde V, Tomasevic L, Akopian I, Stanek K, Saturnino GB, Thielscher A, Bergmann TO, Siebner HR (2019) The non-transcranial TMS-evoked potential is an inherent source of ambiguity in TMS-EEG studies. NeuroImage 185:300–312.

Connors BW, Malenka RC, Silva LR (1988) Two inhibitory postsynaptic potentials, and GABAA and GABAB receptor-mediated responses in neocortex of rat and cat. The Journal of physiology 406:443–468.

Cosmo C, Ferreira C, Miranda JG, do Rosário R, Baptista A, Montoya P, de Sena E (2015) Spreading effect of tDCS in individuals with attention-deficit/hyperactivity disorder as shown by functional cortical networks: a randomized, double-blind, sham-controlled trial. Frontiers in psychiatry 6.

Dale AM, Fischl B, Sereno MI (1999) Cortical surface-based analysis. I. Segmentation and surface reconstruction. NeuroImage 9:179–194.

Darmani G, Ziemann U (2019) Pharmacophysiology of TMS-evoked EEG potentials: A mini-review. Brain stimulation 12:829–831.

Dedoncker J, Brunoni AR, Baeken C, Vanderhasselt M-A (2016) A Systematic Review and Meta-Analysis of the Effects of Transcranial Direct Current Stimulation (tDCS) Over the Dorsolateral Prefrontal Cortex in Healthy and Neuropsychiatric Samples: Influence of Stimulation Parameters. Brain stimulation 9:501–517.

Deng ZD, Lisanby SH, Peterchev AV (2013) Electric field depth-focality tradeoff in transcranial magnetic stimulation: simulation comparison of 50 coil designs. Brain stimulation 6:1–13.

Dieckhöfer A, Waberski TD, Nitsche M, Paulus W, Buchner H, Gobbelé R (2006) Transcranial direct current stimulation applied over the somatosensory cortex – Differential effect on low and high frequency SEPs. Clinical Neurophysiology 117:2221–2227.

Du X, Choa FS, Summerfelt A, Rowland LM, Chiappelli J, Kochunov P, Hong LE (2017) N100 as a generic cortical electrophysiological marker based on decomposition of TMS-evoked potentials across five anatomic locations. Experimental brain research 235:69–81.

Faria P, Hallett M, Miranda PC (2011) A finite element analysis of the effect of electrode area and inter- electrode distance on the spatial distribution of the current density in tDCS. Journal of neural engineering 8:066017–066017.

Faul F, Erdfelder E, Buchner A, Lang A-G (2009) Statistical power analyses using G*Power 3.1: Tests for correlation and regression analyses. Behavior Research Methods 41:1149–1160.

Fecchio M, Pigorini A, Comanducci A, Sarasso S, Casarotto S, Premoli I, Derchi C-C, Mazza A, Russo S, Resta F, Ferrarelli F, Mariotti M, Ziemann U, Massimini M, Rosanova M (2017) The spectral features of EEG responses to transcranial magnetic stimulation of the primary motor cortex depend on the amplitude of the motor evoked potentials. PLoS One 12:e0184910–e0184910.

Ferreri F, Pasqualetti P, Maatta S, Ponzo D, Ferrarelli F, Tononi G, Mervaala E, Miniussi C, Rossini PM (2011) Human brain connectivity during single and paired pulse transcranial magnetic stimulation. NeuroImage 54:90–102.

Fitzgerald PB (2021) Targeting repetitive transcranial magnetic stimulation in depression: do we really know what we are stimulating and how best to do it? Brain stimulation 14:730–736.

Gaetz W, Edgar JC, Wang DJ, Roberts TP (2011) Relating MEG measured motor cortical oscillations to resting γ-aminobutyric acid (GABA) concentration. NeuroImage 55:616–621.

Gaspar P, Bloch B, Le Moine C (1995) D1 and D2 receptor gene expression in the rat frontal cortex: cellular localization in different classes of efferent neurons. The European journal of neuroscience 7:1050–1063.

Ghasemian-Shirvan E, Mosayebi-Samani M, Farnad L, Kuo M-F, Meesen RLJ, Nitsche MA (2022) Age- dependent non-linear neuroplastic effects of cathodal tDCS in the elderly population: a titration study. Brain stimulation 15:296–305.

Ghasemian-Shirvan E, Farnad L, Mosayebi-Samani M, Verstraelen S, Meesen RLJ, Kuo M-F, Nitsche MA (2020) Age-related differences of motor cortex plasticity in adults: A transcranial direct current stimulation study. Brain stimulation 13:1588–1599.

Gordon PC, Desideri D, Belardinelli P, Zrenner C, Ziemann U (2018a) Comparison of cortical EEG responses to realistic sham versus real TMS of human motor cortex. Brain stimulation 11:1322–1330.

Gordon PC, Zrenner C, Desideri D, Belardinelli P, Zrenner B, Brunoni AR, Ziemann U (2018b) Modulation of cortical responses by transcranial direct current stimulation of dorsolateral prefrontal cortex: A resting-state EEG and TMS-EEG study. Brain stimulation 11:1024–1032.

Gramfort A, Papadopoulo T, Olivi E, Clerc M (2010) OpenMEEG: opensource software for quasistatic bioelectromagnetics. BioMedical Engineering OnLine 9:45.

Hernandez-Pavon JC, Metsomaa J, Mutanen T, Stenroos M, Maki H, Ilmoniemi RJ, Sarvas J (2012) Uncovering neural independent components from highly artifactual TMS-evoked EEG data. Journal of neuroscience methods 209:144–157.

Herwig U, Satrapi P, Schonfeldt-Lecuona C (2003) Using the international 10-20 EEG system for positioning of transcranial magnetic stimulation. Brain Topogr 16:95–99.

Hill AT, Rogasch NC, Fitzgerald PB, Hoy KE (2017) Effects of prefrontal bipolar and high-definition transcranial direct current stimulation on cortical reactivity and working memory in healthy adults. NeuroImage 152:142–157.

Hill AT, Rogasch NC, Fitzgerald PB, Hoy KE (2019) Impact of concurrent task performance on transcranial direct current stimulation (tDCS)-Induced changes in cortical physiology and working memory. Cortex 113:37–57.

Howard MW, Rizzuto DS, Caplan JB, Madsen JR, Lisman J, Aschenbrenner-Scheibe R, Schulze-Bonhage A, Kahana MJ (2003) Gamma oscillations correlate with working memory load in humans. Cerebral cortex (New York, NY : 1991) 13:1369-1374.

Hyvärinen A, Oja E (2000) Independent component analysis: algorithms and applications. Neural Networks 13:411–430.

Ilmoniemi RJ, Kicic D (2010) Methodology for combined TMS and EEG. Brain Topogr 22:233–248.

Ilmoniemi RJ, Virtanen J, Ruohonen J, Karhu J, Aronen HJ, Näätänen R, Katila T (1997) Neuronal responses to magnetic stimulation reveal cortical reactivity and connectivity. Neuroreport 8:3537–3540.

Jamil A, Batsikadze G, Kuo HI, Labruna L, Hasan A, Paulus W, Nitsche MA (2017) Systematic evaluation of the impact of stimulation intensity on neuroplastic after-effects induced by transcranial direct current stimulation. The Journal of physiology 595:1273–1288.

Karuza EA, Balewski ZZ, Hamilton RH, Medaglia JD, Tardiff N, Thompson-Schill SL (2016) Mapping the Parameter Space of tDCS and Cognitive Control via Manipulation of Current Polarity and Intensity. Front Hum Neurosci 10.

Keeser D, Meindl T, Bor J, Palm U, Pogarell O, Mulert C, Brunelin J, Moller HJ, Reiser M, Padberg F (2011) Prefrontal transcranial direct current stimulation changes connectivity of resting-state networks during fMRI. The Journal of neuroscience : the official journal of the Society for Neuroscience 31:15284–15293.

Komssi S, Kahkonen S (2006) The novelty value of the combined use of electroencephalography and transcranial magnetic stimulation for neuroscience research. Brain research reviews 52:183–192.

König F, Belardinelli P, Liang C, Desideri D, Müller-Dahlhaus F, Gordon PC, Zipser C, Zrenner C, Ziemann U (2019) TMS-EEG signatures of glutamatergic neurotransmission in human cortex. bioRxiv:555920.

Kozel FA, Nahas Z, deBrux C, Molloy M, Lorberbaum JP, Bohning D, Risch SC, George MS (2000) How coil- cortex distance relates to age, motor threshold, and antidepressant response to repetitive transcranial magnetic stimulation. The Journal of neuropsychiatry and clinical neurosciences 12:376–384.

Kuo M-F, Chen P-S, Nitsche MA (2017) The application of tDCS for the treatment of psychiatric diseases. International Review of Psychiatry 29:146–167.

Kuo MF, Paulus W, Nitsche MA (2008) Boosting focally-induced brain plasticity by dopamine. Cerebral cortex (New York, NY : 1991) 18:648-651.

Laakso I, Tanaka S, Koyama S, De Santis V, Hirata A (2015) Inter-subject Variability in Electric Fields of Motor Cortical tDCS. Brain stimulation 8:906–913.

Laakso I, Mikkonen M, Koyama S, Hirata A, Tanaka S (2019) Can electric fields explain inter-individual variability in transcranial direct current stimulation of the motor cortex? Scientific reports 9:626.

Lang N, Siebner HR, Ward NS, Lee L, Nitsche MA, Paulus W, Rothwell JC, Lemon RN, Frackowiak RS (2005) How does transcranial DC stimulation of the primary motor cortex alter regional neuronal activity in the human brain? The European journal of neuroscience 22:495–504.

Lefaucheur JP et al. (2017) Evidence-based guidelines on the therapeutic use of transcranial direct current stimulation (tDCS). Clinical neurophysiology : official journal of the International Federation of Clinical Neurophysiology 128:56–92.

Lioumis P, Kicic D, Savolainen P, Makela JP, Kahkonen S (2009) Reproducibility of TMS-Evoked EEG responses. Human brain mapping 30:1387–1396.

Maris E, Oostenveld R (2007) Nonparametric statistical testing of EEG- and MEG-data. Journal of neuroscience methods 164:177–190.

Matsunaga K, Nitsche MA, Tsuji S, Rothwell JC (2004) Effect of transcranial DC sensorimotor cortex stimulation on somatosensory evoked potentials in humans. Clinical neurophysiology : official journal of the International Federation of Clinical Neurophysiology 115:456–460.

McConnell KA, Nahas Z, Shastri A, Lorberbaum JP, Kozel FA, Bohning DE, George MS (2001) The transcranial magnetic stimulation motor threshold depends on the distance from coil to underlying cortex: a replication in healthy adults comparing two methods of assessing the distance to cortex. Biological psychiatry 49:454–459.

McFadden JL, Borckardt JJ, George MS, Beam W (2011) Reducing procedural pain and discomfort associated with transcranial direct current stimulation. Brain stimulation 4:38–42.

Miranda PC, Mekonnen A, Salvador R, Ruffini G (2013) The electric field in the cortex during transcranial current stimulation. NeuroImage 70:48–58.

Mosayebi-Samani M, Melo L, Agboada D, Nitsche MA, Kuo M-F (2020) Ca2+ channel dynamics explain the nonlinear neuroplasticity induction by cathodal transcranial direct current stimulation over the primary motor cortex. European Neuropsychopharmacology 38:63–72.

Mosayebi-Samani M, Jamil A, Salvador R, Ruffini G, Haueisen J, Nitsche MA (2021) The impact of individual electrical fields and anatomical factors on the neurophysiological outcomes of tDCS: A TMS-MEP and MRI study. Brain stimulation 14:316–326.

Mosayebi Samani M, Agboada D, Jamil A, Kuo M-F, Nitsche MA (2019) Titrating the neuroplastic effects of cathodal transcranial direct current stimulation (tDCS) over the primary motor cortex. Cortex 119:350–361.

Mutanen T, Maki H, Ilmoniemi RJ (2013) The effect of stimulus parameters on TMS-EEG muscle artifacts. Brain stimulation 6:371–376.

Mutanen TP, Kukkonen M, Nieminen JO, Stenroos M, Sarvas J, Ilmoniemi RJ (2016) Recovering TMS- evoked EEG responses masked by muscle artifacts. NeuroImage 139:157–166.

Nikouline V, Ruohonen J, Ilmoniemi RJ (1999) The role of the coil click in TMS assessed with simultaneous EEG. Clinical neurophysiology : official journal of the International Federation of Clinical Neurophysiology 110:1325–1328.

Nikulin VV, Kicić D, Kähkönen S, Ilmoniemi RJ (2003) Modulation of electroencephalographic responses to transcranial magnetic stimulation: evidence for changes in cortical excitability related to movement. The European journal of neuroscience 18:1206–1212.

Nitsche M, Koschack J, Pohlers H, Hullemann S, Paulus W, Happe S (2012) Effects of Frontal Transcranial Direct Current Stimulation on Emotional State and Processing in Healthy Humans. Frontiers in psychiatry 3.

Nitsche M, Fricke K, Henschke U, Schlitterlau A, Liebetanz D, Lang N, Henning S, Tergau F, Paulus W (2003) Pharmacological modulation of cortical excitability shifts induced by transcranial direct current stimulation in humans. The Journal of physiology 553:293–301.

Nitsche MA, Liebetanz D, Schlitterlau A, Henschke U, Fricke K, Frommann K, Lang N, Henning S, Paulus W, Tergau F (2004) GABAergic modulation of DC stimulation-induced motor cortex excitability shifts in humans. European Journal of Neuroscience 19:2720–2726.

Nitsche MA, Cohen LG, Wassermann EM, Priori A, Lang N, Antal A, Paulus W, Hummel F, Boggio PS, Fregni F (2008) Transcranial direct current stimulation: state of the art 2008. Brain stimulation 1:206–223.

Oldfield RC (1971) The assessment and analysis of handedness: the Edinburgh inventory. Neuropsychologia 9:97–113.

Oostenveld R, Fries P, Maris E, Schoffelen J-M (2011) FieldTrip: open source software for advanced analysis of MEG, EEG, and invasive electrophysiological data. Computational intelligence and neuroscience 2011.

Opitz A, Paulus W, Will S, Antunes A, Thielscher A (2015) Determinants of the electric field during transcranial direct current stimulation. NeuroImage 109:140–150.

Pellicciari MC, Brignani D, Miniussi C (2013) Excitability modulation of the motor system induced by transcranial direct current stimulation: a multimodal approach. NeuroImage 83:569–580.

Polanía R, Paulus W, Antal A, Nitsche MA (2011) Introducing graph theory to track for neuroplastic alterations in the resting human brain: a transcranial direct current stimulation study. NeuroImage 54:2287–2296.

Poreisz C, Boros K, Antal A, Paulus W (2007) Safety aspects of transcranial direct current stimulation concerning healthy subjects and patients. Brain research bulletin 72:208–214.

Premoli I, Rivolta D, Espenhahn S, Castellanos N, Belardinelli P, Ziemann U, Müller-Dahlhaus F (2014a) Characterization of GABAB-receptor mediated neurotransmission in the human cortex by paired- pulse TMS-EEG. NeuroImage 103:152–162.

Premoli I, Castellanos N, Rivolta D, Belardinelli P, Bajo R, Zipser C, Espenhahn S, Heidegger T, Müller- Dahlhaus F, Ziemann U (2014b) TMS-EEG Signatures of GABAergic Neurotransmission in the Human Cortex. The Journal of Neuroscience 34:5603.

Purpura DP, McMurtry JG (1965) Intracellular activities and evoked potential changes during polarization of motor cortex. Journal of neurophysiology 28:166–185.

Rocchi L, Di Santo A, Brown K, Ibáñez J, Casula E, Rawji V, Di Lazzaro V, Koch G, Rothwell J (2021) Disentangling EEG responses to TMS due to cortical and peripheral activations. Brain stimulation 14:4–18.

Rogasch NC, Fitzgerald PB (2013) Assessing cortical network properties using TMS-EEG. Human brain mapping 34:1652–1669.

Rogasch NC, Thomson RH, Farzan F, Fitzgibbon BM, Bailey NW, Hernandez-Pavon JC, Daskalakis ZJ, Fitzgerald PB (2014) Removing artefacts from TMS-EEG recordings using independent component analysis: Importance for assessing prefrontal and motor cortex network properties. NeuroImage 101:425–439.

Rogasch NC, Sullivan C, Thomson RH, Rose NS, Bailey NW, Fitzgerald PB, Farzan F, Hernandez-Pavon JC (2017) Analysing concurrent transcranial magnetic stimulation and electroencephalographic data: A review and introduction to the open-source TESA software. NeuroImage 147:934–951.

Rogasch NC, Zipser C, Darmani G, Mutanen TP, Biabani M, Zrenner C, Desideri D, Belardinelli P, Müller- Dahlhaus F, Ziemann U (2020) The effects of NMDA receptor blockade on TMS-evoked EEG potentials from prefrontal and parietal cortex. Scientific reports 10:3168.

Romero Lauro LJ, Rosanova M, Mattavelli G, Convento S, Pisoni A, Opitz A, Bolognini N, Vallar G (2014) TDCS increases cortical excitability: direct evidence from TMS-EEG. Cortex 58:99–111.

Rossi S, Hallett M, Rossini PM, Pascual-Leone A, Group SoTC (2009) Safety, ethical considerations, and application guidelines for the use of transcranial magnetic stimulation in clinical practice and research. Clinical Neurophysiology 120:2008–2039.

Rusjan PM, Barr MS, Farzan F, Arenovich T, Maller JJ, Fitzgerald PB, Daskalakis ZJ (2010) Optimal transcranial magnetic stimulation coil placement for targeting the dorsolateral prefrontal cortex using novel magnetic resonance image-guided neuronavigation. Human brain mapping 31:1643–1652.

Stagg CJ, Best JG, Stephenson MC, O’Shea J, Wylezinska M, Kincses ZT, Morris PG, Matthews PM, Johansen-Berg H (2009) Polarity-sensitive modulation of cortical neurotransmitters by transcranial stimulation. Journal of Neuroscience 29:5202–5206.

Sun Y, Dhamne SC, Carretero-Guillén A, Salvador R, Goldenberg MC, Godlewski BR, Pascual-Leone A, Madsen JR, Stone SSD, Ruffini G, Márquez-Ruiz J, Rotenberg A (2020) Drug-Responsive Inhomogeneous Cortical Modulation by Direct Current Stimulation. Annals of neurology 88:489–502.

Tadel F, Baillet S, Mosher JC, Pantazis D, Leahy RM (2011) Brainstorm: a user-friendly application for MEG/EEG analysis. Comput Intell Neurosci 2011:879716.

ter Braack EM, de Vos CC, van Putten MJ (2015) Masking the Auditory Evoked Potential in TMS-EEG: A Comparison of Various Methods. Brain Topogr 28:520–528.

Thielscher A, Antunes A, Saturnino GB (2015) Field modeling for transcranial magnetic stimulation: A useful tool to understand the physiological effects of TMS? Conference proceedings : Annual International Conference of the IEEE Engineering in Medicine and Biology Society IEEE Engineering in Medicine and Biology Society Annual Conference 2015:222–225.

Tremblay S et al. (2019) Clinical utility and prospective of TMS–EEG. Clinical Neurophysiology 130:802–844.

van den Broek SP, Reinders F, Donderwinkel M, Peters MJ (1998) Volume conduction effects in EEG and MEG. Electroencephalography and Clinical Neurophysiology 106:522–534.

Voineskos AN, Farzan F, Barr MS, Lobaugh NJ, Mulsant BH, Chen R, Fitzgerald PB, Daskalakis ZJ (2010) The role of the corpus callosum in transcranial magnetic stimulation induced interhemispheric signal propagation. Biological psychiatry 68:825–831.

Womelsdorf T, Schoffelen JM, Oostenveld R, Singer W, Desimone R, Engel AK, Fries P (2007) Modulation of neuronal interactions through neuronal synchronization. Science 316:1609–1612.

Yavari F, Jamil A, Mosayebi Samani M, Vidor LP, Nitsche MA (2018) Basic and functional effects of transcranial Electrical Stimulation (tES)—An introduction. Neuroscience & Biobehavioral Reviews 85:81–92.

Zhang Y, Chen Y, Bressler SL, Ding M (2008) Response preparation and inhibition: the role of the cortical sensorimotor beta rhythm. Neuroscience 156:238–246.

