## Supplemental TablesFigures for "Transferability of cathodal tDCS effects from the primary motor to the dorsolateral prefrontal cortex: a multimodal TMS-EEG study"

**Table 1-1- TEPs and MEPs included in the data.** MEP amplitudes were first visually inspected to exclude MEP trials: 1) in which background electromyographic activity was present, and 2) with bad TEP trials. Therefore, an identical number of TEPs and MEPs was included for statistical analysis. In addition, the bad EEG channels were interpolated after preprocessing the EEG raw data. Furthermore, an independent component analysis (ICA) was used to remove artifacts of the recorded TEPs. Data are presented as mean ± standard deviation (SD).

| | | | Total trials | Interpolated EEG Channels | Removed MEPs | removed ICA components |
|---|---|---|---|---|---|---|
| **Motor Cortex** | **Sham** | **Baseline** | 105 ± 11 | 1.16 ± 0.98 | 3 ± 4.6 | 6 ±2.3 |
| | | **0 min** | 103 ± 11 | 1.05 ± 1.05 | 5 ± 4.3 | 7 ± 3.1 |
| | | **30 min** | 101 ± 12 | 1.33 ± 0.90 | 3 ± 5.2 | 5 ± 3.5 |
| | | **60 min** | 102 ± 11 | 1.27 ± 1.1 | 5 ± 3.4 | 6 ± 3.2 |
| | | **120min** | 111 ± 7 | 0.94 ± 0.87 | 4 ± 3.2 | 5 ± 4.4 |
| | **Low dosage** | **Baseline** | 102 ± 12 | 1.27 ± 1.07 | 3 ± 3.2 | 4 ± 4.2 |
| | | **0 min** | 103 ± 10 | 1.05 ± 0.72 | 4 ± 3.2 | 6 ± 4.3 |
| | | **30 min** | 100 ± 10 | 1.27 ± 1.17 | 6 ± 2.2 | 5 ± 5.3 |
| | | **60 min** | 104 ± 10 | 1.11 ± 1.02 | 5 ± 3.3 | 6 ± 3.6 |
| | | **120min** | 102 ± 12 | 1.72 ± 0.89 | 4 ± 4.3 | 4 ± 3.8 |
| | **Medium Dosage** | **Baseline** | 100 ± 11 | 1.16 ± 1.15 | 5 ± 5.3 | 4 ± 4.6 |
| | | **0 min** | 103 ± 13 | 1.66 ± 1.23 | 4 ± 2.5 | 5 ± 4.4 |
| | | **30 min** | 92 ± 11 | 0.94 ± 1.01 | 4 ± 2.8 | 6 ± 3.2 |
| | | **60 min** | 102 ± 13 | 1.55 ± 0.98 | 5 ± 3.6 | 7 ± 4.1 |
| | | **120min** | 101 ± 14 | 1.05 ± 0.93 | 5 ± 5.1 | 8 ± 2.2 |
| | **High Dosage** | **Baseline** | 93 ± 11 | 1.44 ± 1.09 | 3 ± 2.1 | 5 ± 3.7 |
| | | **0 min** | 96 ± 11 | 1.11 ± 0.83 | 4 ± 5.3 | 6 ± 4.6 |
| | | **30 min** | 105 ± 8 | 1.27 ± 1.17 | 5 ± 4.2 | 4 ± 5.9 |
| | | **60 min** | 103 ± 10 | 1.33 ± 1.18 | 3 ± 2.4 | 4 ± 3.6 |

| | | | | | | |
|---|---|---|---|---|---|---|
| | | **120min** | 106 ± 10 | 1.33 ± 1.23 | 4 ± 5.6 | 5 ± 4.8 |
| **Prefrontal Cortex** | **Sham** | **Baseline** | 103 ± 8 | 1.64 ± 0.99 | ----- | 6 ± 2.6 |
|  |  | **0 min** | 103 ± 12 | 1.82 ± 1.07 | ----- | 5 ± 3.3 |
|  |  | **30 min** | 103 ± 15 | 1.47 ± 1.06 | ----- | 4 ± 3.8 |
|  |  | **60 min** | 98 ± 11 | 1.82 ± 1.13 | ----- | 6 ± 2.5 |
|  |  | **120min** | 105 ± 11 | 0.88 ± 1.05 | ----- | 5 ± 4.6 |
|  | **Low dosage** | **Baseline** | 94 ± 12 | 1.41 ± 0.87 | ----- | 6 ± 4.2 |
|  |  | **0 min** | 100 ± 11 | 1.29 ± 0.98 | ----- | 5 ± 5.3 |
|  |  | **30 min** | 95 ± 12 | 1.35 ± 1.16 | ----- | 4 ± 2.5 |
|  |  | **60 min** | 100 ± 11 | 1.29 ± 1.26 | ----- | 6 ± 3.6 |
|  |  | **120min** | 95 ± 12 | 1.64 ± 0.86 | ----- | 5 ± 4.7 |
|  | **Medium Dosage** | **Baseline** | 95 ± 15 | 1.47 ± 1.12 | ----- | 6 ± 4.2 |
|  |  | **0 min** | 95 ± 14 | 1.17 ± 0.95 | ----- | 4 ± 4.1 |
|  |  | **30 min** | 95 ± 9 | 1.18 ± 1.07 | ----- | 5 ± 3.3 |
|  |  | **60 min** | 94 ± 13 | 1.05 ± 1.08 | ----- | 6 ± 2.3 |
|  |  | **120min** | 96 ± 11 | 1.05 ± 0.96 | ----- | 7 ± 4.4 |
|  | **High Dosage** | **Baseline** | 94 ± 11 | 1.23 ± 1.03 | ----- | 6 ± 5.6 |
|  |  | **0 min** | 96 ± 13 | 1.41 ± 1.00 | ----- | 4 ± 4.2 |
|  |  | **30 min** | 98 ± 19 | 1.82 ± 1.01 | ----- | 5 ± 3.5 |
|  |  | **60 min** | 99 ± 7 | 1.35 ± 0.93 | ----- | 6 ± 2.2 |
|  |  | **120min** | 93 ± 12 | 1.52 ± 1.06 | ----- | 7 ± 4.3 |

**Table 2-1- Results of the ANOVAs conducted for tDCS-induced TEP alterations (normalized values).** The statistical results indicate tDCS-induced effects for the early (P30 and P60) TEP peaks, with no one-to-one transferability of tDCS effects from the motor to the prefrontal cortex. Asterisks indicate significant effects ($p < .05$), d.f. = degrees of freedom, $\eta_p^2$ = partial eta squared.

| | Factors | d.f., Error | F Value | p Value | $\eta_p^2$ |
|---|---|---|---|---|---|
| **P30** | **Condition** | **3, 30** | **9.610** | **<0.001*** | **0.490** |
| | **Time-point** | **3, 30** | **3.382** | **0.031*** | **0.253** |
| | **Stimulation site** | 1, 10 | 0.568 | 0.469 | 0.054 |
| | **Condition × Time-point** | **9. 90** | **2.381** | **0.018*** | **0.192** |
| | **Condition × Stimulation site** | 1.378, 13.781 | 0.310 | 0.658 | 0.030 |
| | **Time-point × Stimulation site** | 3, 30 | 1.308 | 0.290 | 0.116 |
| | **Condition × Time-point × Stimulation site** | 9, 90 | 1.225 | 0.290 | 0.109 |
| | **Condition** | 3, 15 | 0.294 | 0.829 | 0.056 |

| | | | | | |
|---|---|---|---|---|---|
| **N45** | **Time-point** | 1.262, 6.309 | 1.327 | 0.289 | 0.215 |
| | **Stimulation site** | 1, 5 | 2.444 | 0.179 | 0.328 |
| | **Condition × Time-point** | 9, 45 | 0.720 | 0.688 | 0.126 |
| | **Condition × Stimulation site** | 3, 15 | 1.372 | 0.289 | 0.215 |
| | **Time-point × Stimulation site** | 3, 15 | 1.055 | 0.398 | 0.174 |
| | **Condition × Time-point × Stimulation site** | 9, 45 | 1.626 | 0.137 | 0.245 |
| **P60** | **Condition** | **3, 15** | **3.441** | **0.034*** | **0.408** |
| | **Time-point** | 3, 20 | 1.054 | 0.405 | 0.174 |
| | **Stimulation site** | 1, 5 | 0.001 | 0.994 | 0.002 |
| | **Condition × Time-point** | 4.314, 51.763 | 0.431 | 0.799 | 0.035 |
| | **Condition × Stimulation site** | 3, 15 | 0.560 | 0.740 | 0.138 |
| | **Time-point × Stimulation site** | 4, 20 | 0.158 | 0.957 | 0.031 |
| | **Condition × Time-point × Stimulation site** | **4.648, 55.777** | **2.185** | **0.035*** | **0.122** |
| **N100** | **Condition** | 3, 42 | 2.518 | 0.071 | 0.152 |
| | **Time-point** | 3, 42 | 1.080 | 0.368 | 0.072 |
| | **Stimulation site** | 1, 14 | 0.611 | 0.447 | 0.042 |
| | **Condition × Time-point** | 9, 126 | 1.863 | 0.063 | 0.117 |
| | **Condition × Stimulation site** | 3, 42 | 0.931 | 0.434 | 0.062 |
| | **Time-point × Stimulation site** | 1.935, 27.095 | 1.568 | 0.227 | 0.101 |
| | **Condition × Time-point × Stimulation site** | 9, 126 | 1.147 | 0.335 | 0.076 |
| **ΔP200** | **Condition** | 3, 36 | 0.191 | 0.902 | 0.016 |
| | **Time-point** | 3, 36 | 2.432 | 0.082 | 0.168 |
| | **Stimulation site** | 1, 12 | 0.245 | 0.629 | 0.020 |
| | **Condition × Time-point** | 9, 108 | 0.770 | 0.645 | 0.060 |
| | **Condition × Stimulation site** | 3, 36 | 1.615 | 0.203 | 0.119 |
| | **Time-point × Stimulation site** | 3, 36 | 2.114 | 0.116 | 0.150 |
| | **Condition × Time-point × Stimulation site** | 3.611, 43.336 | 1.293 | 0.249 | 0.097 |

**Table 3-1- Results of the ANOVAs conducted for tDCS-induced alterations of cortical oscillations (normalized values).** The secondary rmANOVAs conducted to test the effects of active tDCS conditions on TMS-evoked oscillations vs. sham showed no significant effect of neither the main factors condition, time-point, and stimulation sites, nor their respective interactions. d.f. = degrees of freedom, $\eta_p^2$ = partial eta squared.

| | | **Factors** | **d.f.** | **F Value** | **p Value** | $\boldsymbol{\eta_p^2}$ |
|---|---|---|---|---|---|---|
| **Frequency bands** | **Δθ** | **Condition** | 1.789, 19.680 | 2.318 | 0.129 | 0.174 |
| | | **Time-point** | 3, 33 | 0.461 | 0.712 | 0.040 |
| | | **Stimulation site** | 1, 12 | 4.016 | 0.067 | 0.213 |
| | | **Condition × Time-point** | 3.610, 39.710 | 2.559 | 0.138 | 0.189 |
| | | **Condition × Stimulation site** | 3, 33 | 0.971 | 0.418 | 0.081 |
| | | **Time-point × Stimulation site** | 3, 33 | 0.174 | 0.913 | 0.016 |
| | | **Condition × Time-point × Stimulation site** | 4.297, 47.269 | 0.560 | 0.827 | 0.048 |
| | **Δα** | **Condition** | 3, 24 | 0.334 | 0.801 | 0.040 |
| | | **Time-point** | 3, 24 | 0.792 | 0.510 | 0.090 |
| | | **Stimulation site** | 1, 12 | 2.236 | 0.169 | 0.148 |
| | | **Condition × Time-point** | 2.031,16.252 | 2.139 | 0.149 | 0.211 |
| | | **Condition × Stimulation site** | 3, 24 | 0.790 | 0.511 | 0.090 |
| | | **Time-point × Stimulation site** | 1.290, 10.318 | 0.755 | 0.438 | 0.086 |
| | | **Condition × Time-point × Stimulation site** | 1.495, 11.962 | 1.537 | 0.250 | 0.161 |
| | **Δβ** | **Condition** | 3, 33 | 0.716 | 0.550 | 0.061 |
| | | **Time-point** | 1.664, 18.299 | 0.263 | 0.732 | 0.023 |
| | | **Stimulation site** | 1, 11 | 1.535 | 0.241 | 0.122 |
| | | **Condition × Time-point** | 1.783, 19.617 | 0.429 | 0.634 | 0.038 |
| | | **Condition × Stimulation site** | 3, 33 | 1.223 | 0.317 | 0.100 |
| | | **Time-point × Stimulation site** | 1.316, 14.481 | 1.133 | 0.324 | 0.093 |
| | | **Condition × Time-point × Stimulation site** | 3.315, 29.856 | 0.720 | 0.501 | 0.085 |
| | **Δγ** | **Condition** | 3, 21 | 1.016 | 0.406 | 0.127 |
| | | **Time-point** | 3, 21 | 0.784 | 0.516 | 0.101 |
| | | **Stimulation site** | 1, 7 | 1.570 | 0.250 | 0.183 |
| | | **Condition × Time-point** | 9, 63 | 1.201 | 0.310 | 0.146 |
| | | **Condition × Stimulation site** | 3, 21 | 1.338 | 0.289 | 0.160 |
| | | **Time-point × Stimulation site** | 3, 21 | 0.972 | 0.424 | 0.122 |
| | | **Condition × Time-point × Stimulation site** | 9, 63 | 0.925 | 0.509 | 0.117 |

**Figure 4-2: Results from the statistically significant clusters for the effects of tDCS on TMS-evoked Potentials.** Significant TEP clusters obtained for comparisons between baseline (A) or sham (B) measures and tDCS after-effects across time (POST0, POST30, POST60, and POST120) for each stimulation condition. Significant clusters were only identified for POST0 and/or POST30. There were no significant clusters for the time points POST60, and POST120).

| | Stimulation site | Condition | Time Point | Significant Cluster(s) | Cluster Latency | p-Value | Electrodes |
|---|---|---|---|---|---|---|---|
| A. Comparison vs. Baseline measures | M1 | Low Dosage | POST0 | negative | 23-48ms | 0.024 | Fz, F1, FCz, FC1, FC2, FC4, Cz, C2, C4, CPz, CP1, CP2 |
| | | | POST0 | negative | 170-245ms | 0.029 | POz, PO4, PO3, P5, Oz, O2, O1, PO7, PO8 |
| | | | POST30 | negative | 24-61ms | 0.024 | FCz, Cz, CPz, FC1, FC2, CP1, C2, C4 |
| | | | POST30 | negative | 151-251ms | 0.034 | POz, Oz, PO4, O2, PO3, O1, PO7, P5, P7 |
| | | Medium Dosage | POST30 | positive | 21-54ms | 0.041 | Fz, F1, F3, F2, FCz, FC1, FC2, Cz, CPz, CP1 |
| | | High Dosage | POST0 | negative | 22-61ms | 0.028 | Fz, FCz, Cz, CPz, F1, FC1, FC2, C2, C4, CP1 |
| | | | POST0 | negative | 70-148ms | 0.029 | Pz, POz, Oz, PO4, PO8, PO3, PO7, P5, P7, O1, O2, , |
| | | | POST30 | negative | 21-60ms | 0.024 | Fz, F1, F2, FCz, FC1, FC2, FC4, Cz, C2, C4, Cpz, CP1, CP2 |
| | F3 | Medium Dosage | POST0 | negative | 27-69ms | 0.024 | AFz, Fz, FCz, F2, F4, FC2 |
| | | | POST30 | negative | 62-87ms | 0.036 | AFz, Fz, FCz, FC1, FC2, F2, F4, F6 |
| | | | POST30 | negative | 79-111ms | 0.034 | Fz, FCz, Cz, CPz, F2, F4, FC1, FC2, FC4, C2, C4, C6, C1, C3, CP1, CP2 |
| | | High Dosage | POST0 | negative | 27-72ms | 0.039 | AFz, Fz, FCz, F2, F4, FC2, FC4, FC1, C1 |
| | | | POST30 | negative | 21-60ms | 0.024 | AFz, Fz, FCz, Cz, F2, F4, FC2, FC4, FC6, FC1 |
| | | | POST30 | positive | 148-253ms | 0.026 | Fz, FCz, Cz, F2, FC1, FC2, FC4, FC6, C2, C4, CP2 |

| | | | | | | | |
|---|---|---|---|---|---|---|---|
| **B. Comparison vs. Sham measures** | **M1** | **Medium Dosage** | POST30 | positive | 30-47ms | 0.041 | FCz, Cz, CPz, FC1,CP1, C2, CP2 |
| | | **High Dosage** | POST0 | negative | 24-65ms | 0.032 | Fz, FCz, Cz, F1, F2, F4, FC1, FC2, FC4, C2, C4 |
| | | | POST30 | negative | 31-86ms | 0.021 | FCz, Cz, FC1, FC2, FC4, C2, C4, C6, CP2, CP4 |
| | **F3** | **Medium Dosage** | POST30 | negative | 23-69ms | 0.038 | AFz, Fz, FCz, F1, F2, F4, F6, FC2, FC4, FC6 |
| | | **High Dosage** | POST0 | negative | 34-86ms | 0.042 | AFz, Fz, F2, F4, F6, F8, FC4, FC6, FC8 |
| | | | POST30 | negative | 25-96ms | 0.035 | F2, F4, F6, F8, FC4, FC6, FC8, C6 |

**Figure 8-2: Results from the statistically significant clusters for the effects of tDCS on TMS-evoked Oscillations.** Significant clusters obtained for comparisons between baseline measures and tDCS after-effects on TMS-evoked oscillations across different frequency bands of Theta, Alpha, Beta and Gamma and time (POST0, POST30, POST60, and POST120) for each stimulation condition (low-, medium-, and high-dosage). Significant clusters were only identified for POST0 and/or POST30. There were no significant clusters for the time points POST60, andPOST120. Dashed lines indicate that no significant clusters were identified.

| | | **Condition** | **Frequency Band** | **Significant Cluster(s)** | **Time point** | **Cluster Latency** | **p-Value** | **Electrodes** |
|---|---|---|---|---|---|---|---|---|
| **Comparison vs. Baseline measures** | **M1** | **Low Dosage** | Theta | ----- | ----- | ----- | ----- | ----- |
| | | | Alpha | negative | POST30 | 53-155ms | 0.021 | CPz, Pz, POz, CP1, P1, P3, P5, P2, P4, P6, PO4, PO2, O2 |
| | | | Beta | negative | POST30 | 185-295ms | 0.019 | AFz, Fz, FCz, F2, F4, FC1, FC2, FC4, C2, C4, CP2 |
| | | | Gamma | negative | POST0 | 52-98ms | 0.012 | AFz, Fz, FCz, F2, F4, FC2, FC4, C2, C4 |
| | | | | negative | POST30 | 63-126ms | 0.038 | Fz, FCz, FC1, FC2, FC4, C2, C4, CP2, CP4, CP6 |
| | | **Medium Dosage** | Theta | positive | POST30 | 60-128ms | 0.024 | CPz, Pz, CP1, CP2, CP5 |
|  |  |  | Alpha | ----- | ----- | ----- | ----- | ----- |
| | | | Beta | positive | POST30 | 185-295ms | 0.019 | AFz, Fz, F1, F3, F2 F4, FC2, FC4, FC5 |

| | | | | | | | | |
|---|---|---|---|---|---|---|---|---|
|  |  |  | Gamma | ----- | ----- | ----- | ----- | ----- |
| | | **High Dosage** | Theta | negative | POST0 | 65-118ms | 0.018 | Oz, POz, PO4, O2, P2, P4 |
| | | | | negative | POST30 | 96-185ms | 0.034 | Pz, P4, P6, PO4, Oz, O2 |
| | | | Alpha | negative | POST0 | 59-96ms | 0.018 | Fz, FCz, FC1, FC2, FC4 |
| | | | Beta | negative | POST0 | 54-83ms | 0.032 | Fz, FCz, FC2, FC4, C2, C4 |
| | | | | negative | POST30 | 93-164ms | 0.038 | CP6, P6, P8, PO8, |
| | | | Gamma | negative | POST0 | 87-196ms | 0.022 | Fz, FCz, Cz, F2, FC2, FC4, C2, C4 |
|  | **F3** | **Low Dosage** | Theta | ----- | ----- | ----- | ----- | ----- |
|  |  |  | Alpha | ----- | ----- | ----- | ----- | ----- |
|  |  |  | Beta | ----- | ----- | ----- | ----- | ----- |
|  |  |  | Gamma | ----- | ----- | ----- | ----- | ----- |
| | | **Medium Dosage** | Theta | negative | POST30 | 51-108ms | 0.031 | Pz, POz, P2, O2, P1, P3, P5, P7, TP7, CP1 |
| | | | Alpha | negative | POST30 | 75-213ms | 0.018 | Fpz, AFz, Fz, Fp1, AF7, F2, F4, F6, FT8 |
| | | | Beta | negative | POST0 | 62-93ms | 0.042 | Fz, FCz, F2, F4, FC2, FC4, FC6, C4 |
| | | | | negative | POST30 | 71-131ms | 0.022 | AFz, Fz, FCz, F2, F4, F6, F8, FC1, FC2, FC4, FC6 |
| | | | Gamma | negative | POST30 | 99-288ms | 0.014 | F2, F4, FC2, FC4 |
|  |  | **High Dosage** | Theta | ----- | ----- | ----- | ----- | ----- |
| | | | Alpha | negative | POST0 | 89-201ms | 0.023 | AFz, Fz, F2, F4, F6, F8, FC4, FC6, FT8 |
|  |  |  | Beta | ----- | ----- | ----- | ----- | ----- |
| | | | Gamma | negative | POST0 | 87-196ms | 0.029 | AFz, Fz, FCz, F2, FC2, FC1, C1, C2 |
| | | | | negative | POST30 | 67-116ms | 0.003 | AFz, Fz, FCz, Cz, FC1, FC2, C1, C2, C3 |

**Table 5-1- Frequency table of guessed vs actual received stimulation intensity.** In each session, participants were asked to guess the intensity of tDCS (none, low, medium, high). Note that the study included four tDCS dosages applied over two different stimulation sites. The table contrasts actually applied intensity (rows) with perceived intensity (columns).

| | **Stimulation site** | | **Intensity guessed by participants** | | | |
|---|---|---|---|---|---|---|
| | | | **None** | **Low** | **Medium** | **High** |
| | **Motor Cortex** | **None** | 6 | 7 | 3 | 2 |
| | | **Low** | 6 | 8 | 3 | 1 |

| | | | | | | |
|---|---|---|---|---|---|---|
| **Actual tDCS intensity** | | **Medium** | 3 | 7 | 5 | 3 |
| | | **High** | 1 | 4 | 6 | 7 |
| | **Prefrontal Cortex** | **None** | 6 | 7 | 4 | 1 |
| | | **Low** | 6 | 8 | 4 | 0 |
| | | **Medium** | 2 | 6 | 7 | 3 |
| | | **High** | 2 | 5 | 5 | 6 |

**Table 5-2- Participant ratings of the presence and intensity of side effects.** Visual phenomena, itching, tingling, and pain during stimulation. Skin redness, headache, fatigue, concentration difficulties, nervousness, and sleep problems within 24 hours after stimulation. The presence and intensity of the side effects were rated on a numerical scale ranging from zero to five, zero representing no and five extremely strong sensations. Data are presented as mean ± SD.

| | | **Motor Cortex Stimulation** | | | | **Prefrontal Cortex Stimulation** | | | |
|---|---|---|---|---|---|---|---|---|---|
| | **Side-effects** | **Sham** | **Low Dosage** | **Medium Dosage** | **High Dosage** | **Sham** | **Low Dosage** | **Medium Dosage** | **High Dosage** |
| **During stimulation** | **Visual Phenomenon** | 0.33 ± 0.68 | 0.61 ± 0.69 | 0.61 ± 1.03 | 0.05 ± 0.23 | 0.94 ± 1.05 | 0.88 ± 0.90 | 0.33 ± 0.68 | 0.61 ± 0.69 |
| | **Itching** | 0.33 ± 0.48 | 0.50 ± 0.98 | 0.55 ± 0.92 | 0.22 ± 0.54 | 0.33 ± 0.68 | 0.66 ± 0.69 | 0.33 ± 0.48 | 0.44 ± 0.78 |
| | **Tingling** | 0.66 ± 0.84 | 0.50 ± 0.51 | 0.38 ± 0.84 | 1.00 ± 0.90 | 0.33 ± 0.48 | 0.50 ± 0.98 | 0.66 ± 0.84 | 0.55 ± 0.61 |
| | **Pain** | 0.27 ± 0.46 | 0.38 ± 0.77 | 0.50 ± 0.98 | 0.66 ± 1.08 | 0.66 ± 0.84 | 0.50 ± 0.51 | 0.27 ± 0.46 | 0.38 ± 0.77 |
| **24 hours after stimulation** | **Redness** | 0.27 ± 0.95 | 0.22 ± 0.54 | 0.22 ± 0.54 | 0.44 ± 0.85 | 0.27 ± 0.46 | 0.38 ± 0.77 | 0.27 ± 0.95 | 0.16 ± 0.38 |
| | **Headache** | 0.27 ± 0.46 | 0.27 ± 0.57 | 0.44 ± 0.70 | 0.66 ± 1.13 | 0.27 ± 0.95 | 0.22 ± 0.54 | 0.27 ± 0.46 | 0.27 ± 0.57 |
| | **Fatigue** | 0.66 ± 0.76 | 0.33 ± 0.48 | 0.50 ± 0.85 | 0.72 ± 1.17 | 0.27 ± 0.51 | 0.27 ± 0.57 | 0.66 ± 0.76 | 0.33 ± 0.48 |
| | **Concentration** | 0.33 ± 0.84 | 0.33 ± 0.76 | 0.33 ± 0.84 | 0.38 ± 0.60 | 0.66 ± 0.76 | 0.33 ± 0.48 | 0.33 ± 0.84 | 0.33 ± 0.76 |
| | **Nervousness** | 0.00 ± 0.00 | 0.16 ± 0.51 | 0.05 ± 0.23 | 0.50 ± 0.70 | 0.33 ± 0.84 | 0.33 ± 0.76 | 0.00 ± 0.00 | 0.16 ± 0.51 |
| | **Sleep Problem** | 0.00 ± 0.00 | 0.05 ± 0.23 | 0.11 ± 0.47 | 0.44 ± 0.61 | 0.00 ± 0.00 | 0.16 ± 0.51 | 0.00 ± 0.00 | 0.05 ± 0.23 |

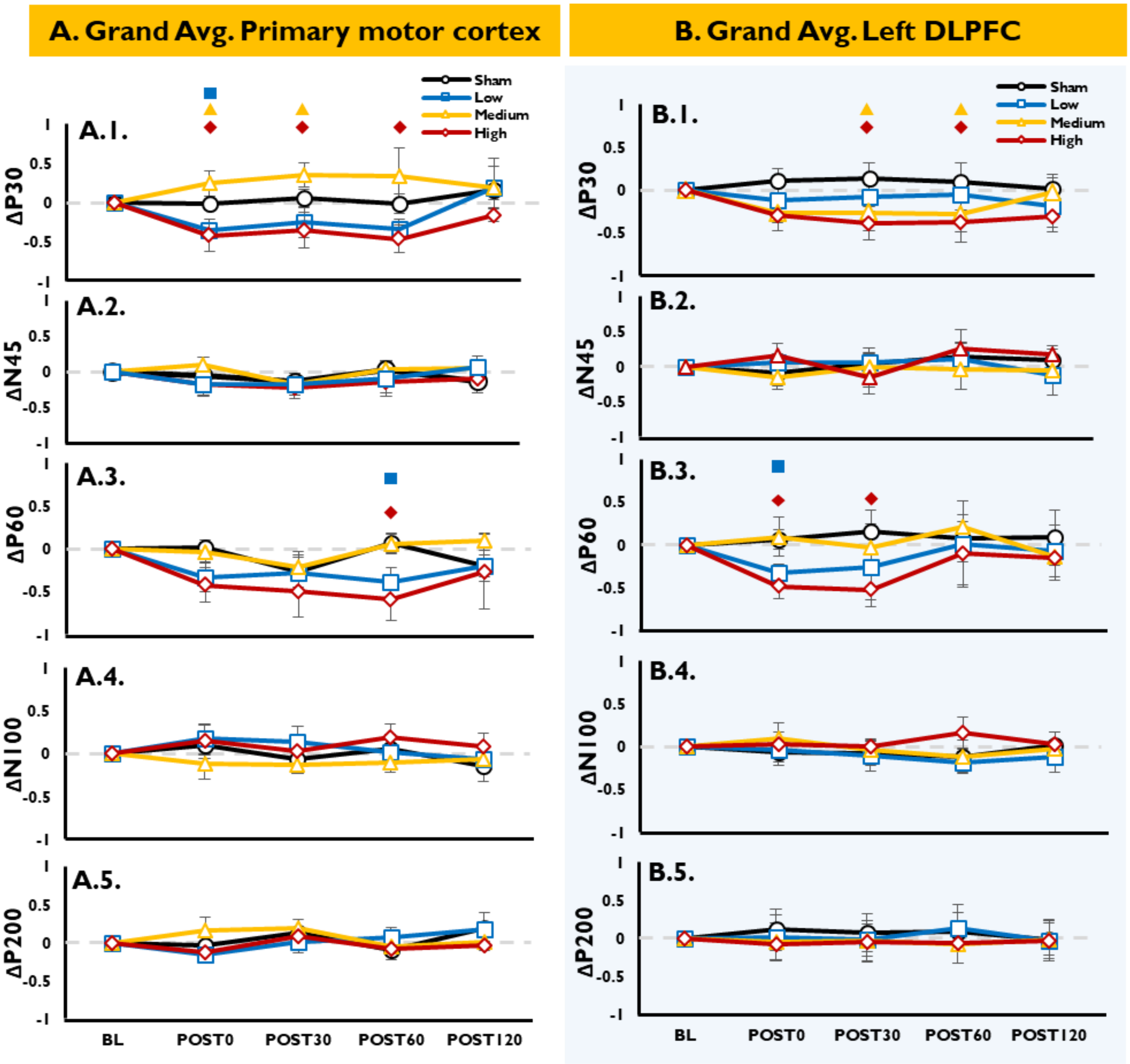


**Figure 3-1. Local effects of tDCS on TMS-evoked Potentials (normalized values).** Low, medium, and high intensities of cathodal tDCS, and sham stimulation were applied over the primary motor (M1) and left dorsolateral prefrontal cortex (PFC). Local tDCS effects were then evaluated, every 30min, from immediately (POST0) to up to two hours after stimulation (POST30, POST60 and POST120), over the $ROI_{M1}$ (averaged FC1 and CP1 electrodes) and $ROI_{PFC}$ (averaged FCz and Fz electrodes). **A1-5, B1-5:** normalized TMS-evoked potentials over M1 and the PFC, respectively. tDCS generated a dosage-dependent, partially non-linear modulation of TEP (ΔP30, ΔN45, ΔP60, ΔN100, ΔP200) over the different stimulation sites, as shown by the amplitude alterations of early ((ΔP30and ΔP60) TEP peaks. Floating symbols show a significant difference of active tDCS conditions (low-dosage■, medium-dosage ▲, and high- dosage◆) vs. sham.

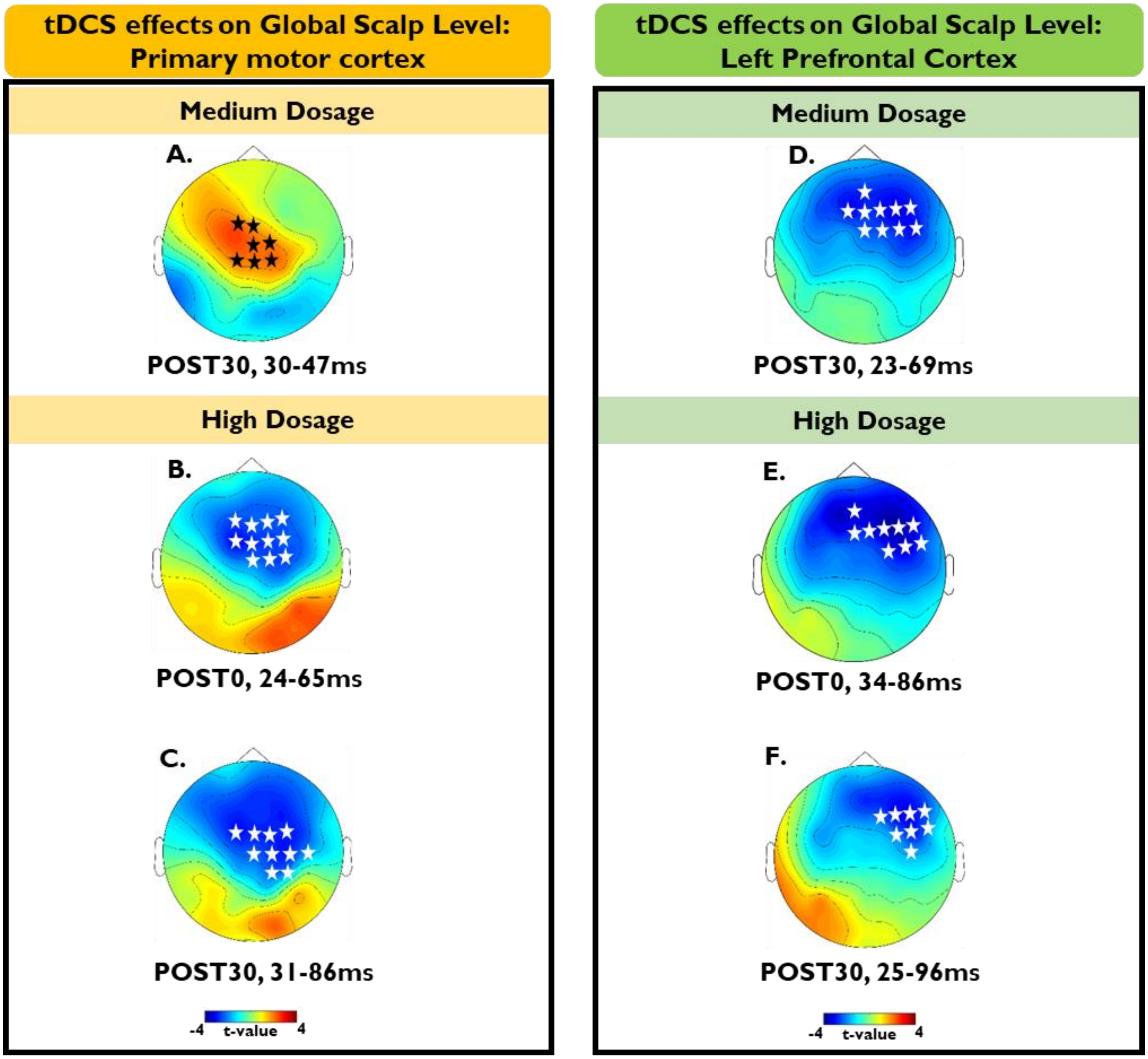


**Figure 4-1. Global effects of tDCS on TMS-evoked Potentials (over M1 and PFC; comparison vs. Sham).** The distributed effects of tDCS were evaluated by cluster-based permutation tests, immediately (POST0) to up to two hours after stimulation (POST30, POST60, POST120), over all electrodes. Topographic plots (distribution of the t-values) shown are only those with significant negative clusters (white stars) or positive clusters (black stars), over M1 (A-C), and PFC (D-F). For further detailed information regarding the specific electrodes forming each cluster please refer to Figure 4-2.

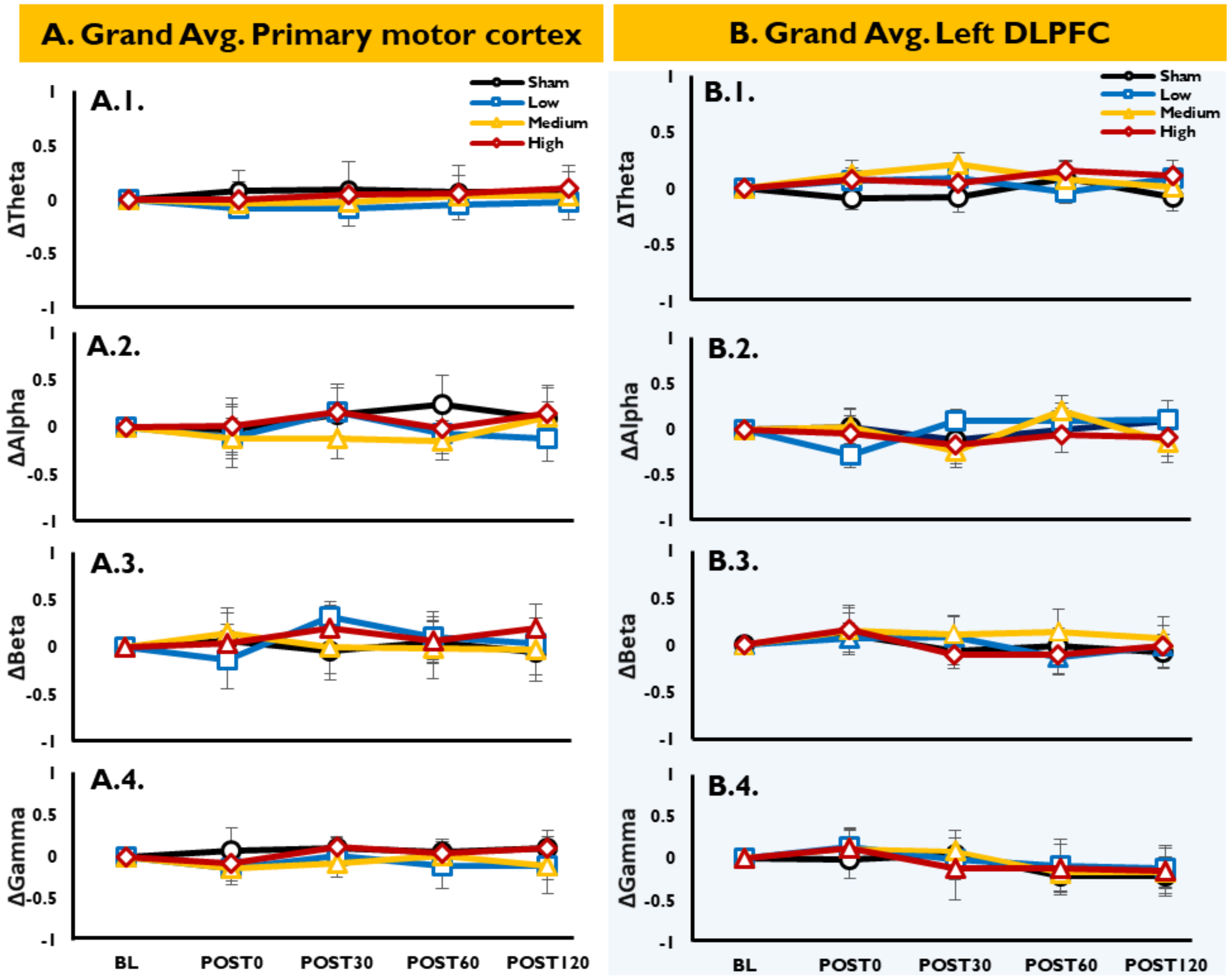


**Figure 7-1. Local effects of tDCS on TMS-evoked Oscillations (normalized values).** Time-frequency representations (TFRs) of oscillatory power were calculated (Morlet wavelet; wavelet width: starting from 2.6 cycles and adding 0.2 cycle for each 1 Hz), and then normalized (db) to the respective baseline (−500 to −100ms), for $ROI_{M1}$ (averaged FC1 and CP1 electrodes) and $ROI_{PFC}$ (averaged FCz and Fz electrodes). Then power estimates were calculated before (BL) and for four time-points (immediately: POST0, 30min: POST30, 60min: POST60 and 120min: POST120) after tDCS, for four separate frequency bands, including Theta (θ; 4-7Hz), Alpha (α; 8-13Hz), Beta (β; 14-29Hz) and Gamma (γ; 30-45Hz), within a time window of 50–300ms. **A.1-4, B.1-4)** Finally, the baseline-normalized values were calculated. Error bars show the standard error of the mean (SEM).

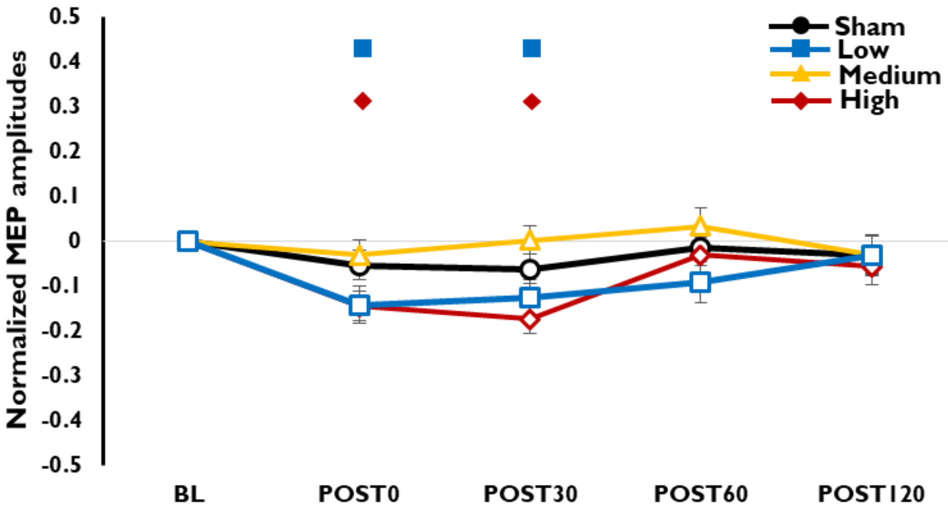


**Figure 10-1. Effects of tDCS on TMS-elicited MEPs (normalized values).** Low, medium, and high cathodal tDCS intensities, and sham stimulation, were applied over the primary motor cortex (M1). The tDCS effects on cortico-spinal excitability were then evaluated immediately (POST0), and for up to two hours after stimulation (POST30, POST60, POST120), using baseline-normalized ΔMEP amplitudes. Floating symbols show a significant difference of active tDCS conditions vs. sham. Error bars show the standard error of the mean (SEM).

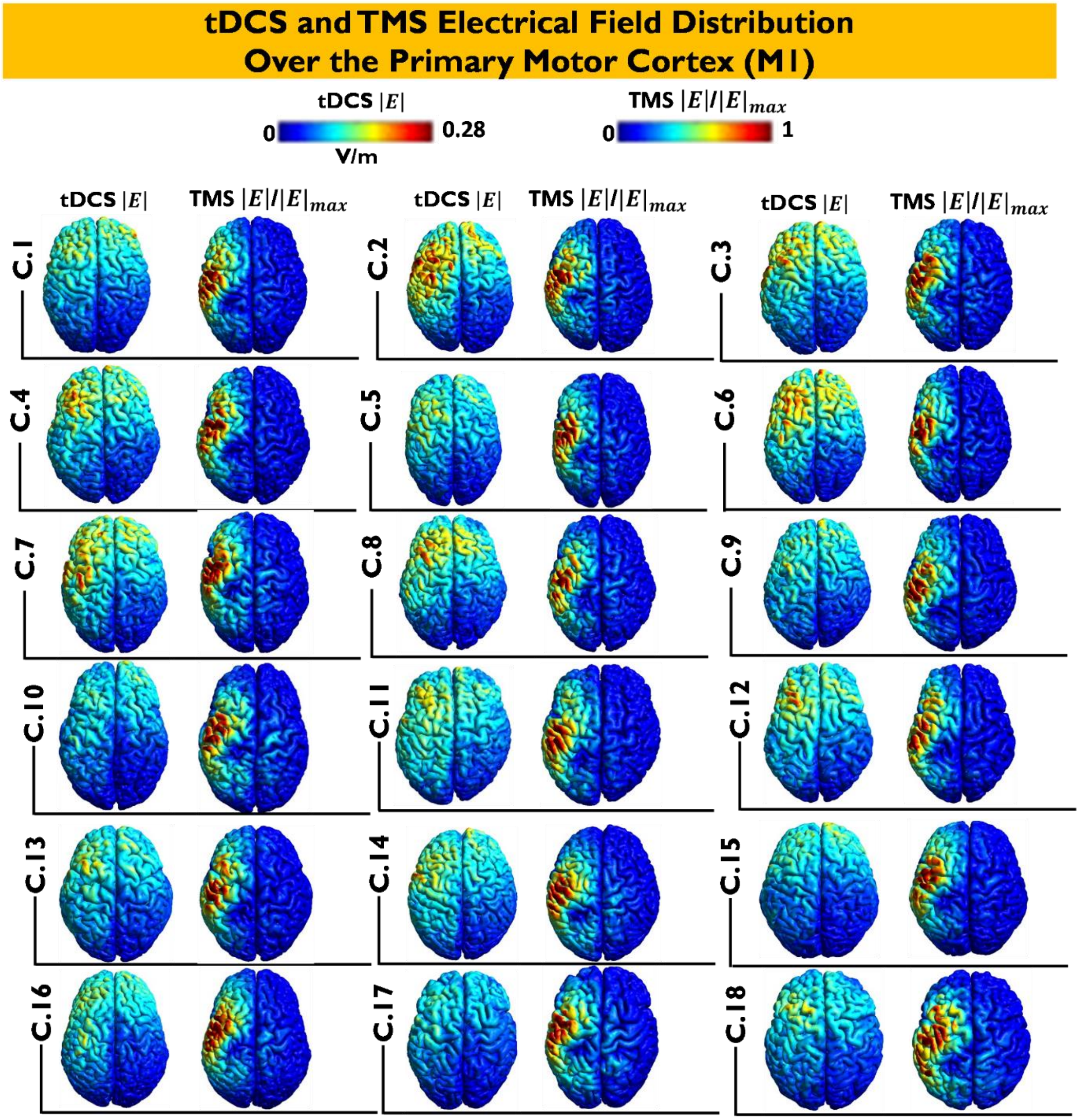


**Figure 11-1. tDCS- and TMS-induced EF over M1.** For each individual, the tDCS- and TMS- induced EFs were calculated. For cathodal tDCS, the target electrode (25cm$^2$) was positioned over the C3 (10-10 EEG standard) position, and the reference electrode (25cm$^2$) was placed contralaterally above the supraorbital area. The resulting electrical fields over the gray matter surface are presented for the tDCS-induced EF magnitude (|E|), and the magnitude of the TMS-induced EF. Note that EFs were calculated for low-intensity tDCS (0.7mA), but, due to the quasi-static assumption, estimated EFs are linearly scaled for other intensities.

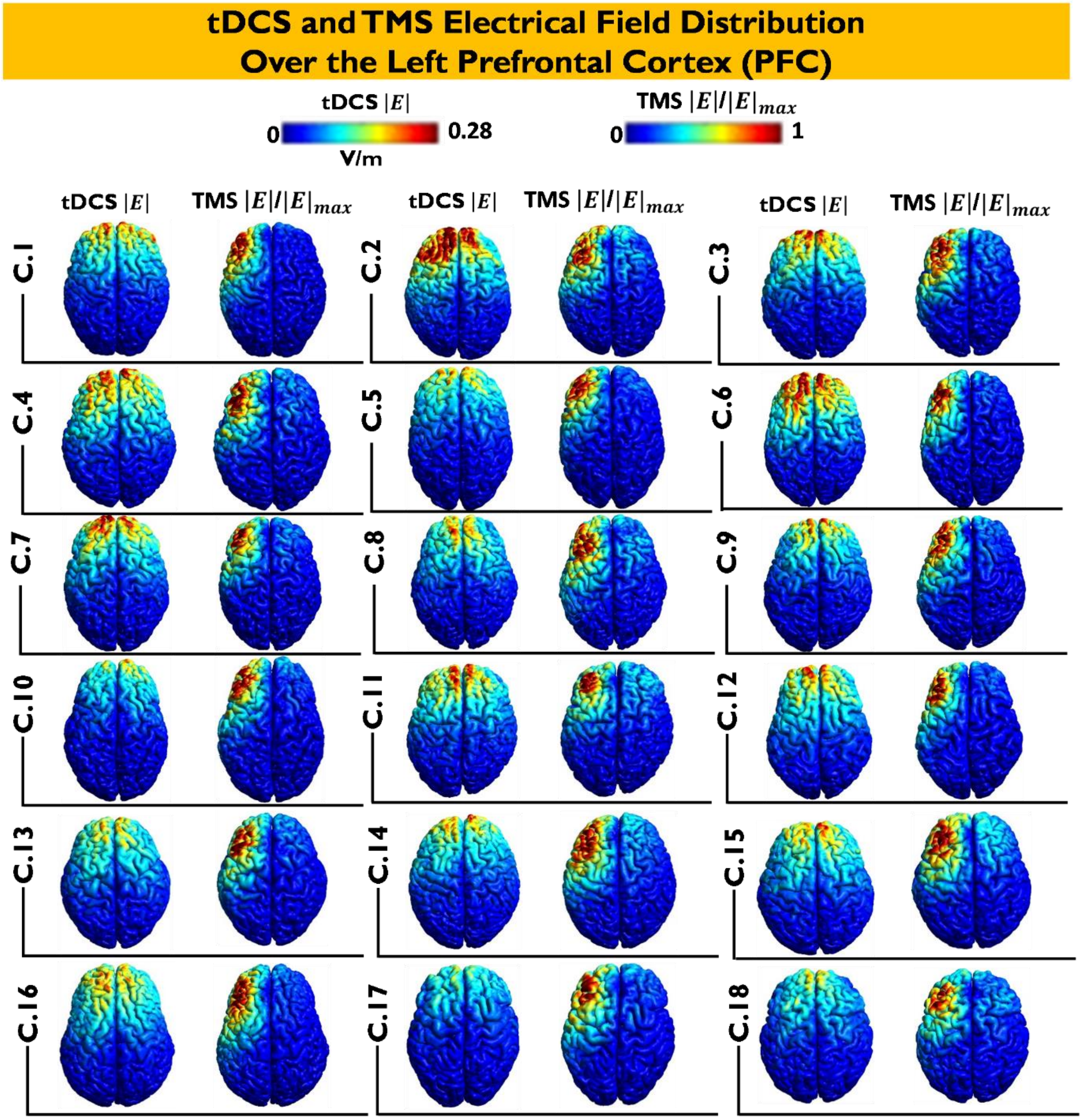


**Figure 11-2. tDCS and TMS-induced EF over the PFC.** For each individual, the tDCS- and TMS-induced EFs were calculated. For cathodal tDCS, the target electrode ($25cm^2$) was positioned over the F3 (10-10 EEG standard) position, and the reference electrode ($25cm^2$) was placed contralaterally above the supraorbital area. The resulting electrical fields over the gray matter surface are presented for the tDCS-induced EF magnitude (|E|), and the magnitude of the TMS-induced EF. Note that EFs were calculated for low-intensity tDCS (0.7mA), but, due to the quasi-static assumption, estimated EFs are linearly scaled for other intensities.
